# Changes in epithelial proportions and transcriptional state underlie major premenopausal breast cancer risks

**DOI:** 10.1101/430611

**Authors:** Lyndsay M. Murrow, Robert J. Weber, Joseph A. Caruso, Christopher S. McGinnis, Kiet Phong, Philippe Gascard, Alexander D. Borowsky, Tejal A. Desai, Matthew Thomson, Thea Tlsty, Zev J. Gartner

## Abstract

The human breast undergoes lifelong remodeling in response to estrogen and progesterone, but hormone exposure also increases breast cancer risk. Here, we use single-cell analysis to identify distinct mechanisms through which breast composition and cell state affect hormone signaling. We show that prior pregnancy reduces the transcriptional response of hormone-responsive (HR+) epithelial cells, whereas high body mass index (BMI) reduces overall HR+ cell proportions. These distinct changes both impact neighboring cells by effectively reducing the magnitude of paracrine signals originating from HR+ cells. Because pregnancy and high BMI are known to protect against hormone-dependent breast cancer in premenopausal women, our findings directly link breast cancer risk with person-to-person heterogeneity in hormone responsiveness. More broadly, our findings illustrate how cell proportions and cell state can collectively impact cell communities through the action of cell-to-cell signaling networks.

## Introduction

The rise and fall of estrogen and progesterone with each menstrual cycle and during pregnancy controls cell growth, survival, and tissue morphology in the human breast. The impact of these changes is profound, and lifetime exposure to cycling hormones is a major modifier of breast cancer risk (*1*). In addition to the dynamics observed within individuals in response to changing hormone levels, there is also a high degree of heterogeneity between individuals in epithelial architecture (*2*), cell composition (*3*), and hormone responsiveness (*4*-*6*), and these differences are thought to impact breast cancer susceptibility. However, because the breast is both highly variable between women and undergoes dynamic changes over time, it has been difficult to link differences in breast cancer risk with specific biological mechanisms in the breast.

While links between cancer risk and specific changes in breast composition, signaling state, or structure are lacking, epidemiological studies have provided clear links between breast cancer risk and several biological variables. Reproductive history and body mass index (BMI) are two such factors that strongly influence breast cancer risk. Pregnancy has two opposing effects: it increases short-term risk by up to 25% (*7*) but decreases lifetime risk by up to 50%, particularly for women with a first pregnancy early in life (*8*). Obesity has opposing effects on risk before versus after menopause: it increases risk in postmenopausal women by around 30% (*9*) but decreases risk in premenopausal women by up to 45% (*10, 11*). The protective effects of both pregnancy against breast cancer and high BMI against premenopausal breast cancer are strongest for estrogen- and progesterone-receptor positive (ER+/PR+) tumors (*11, 12*), suggesting that altered hormone signaling is one mechanism contributing to the protective effect of these two factors. The mechanistic link between pregnancy and the reduction in long-term breast cancer risk remains an open question, but it has been speculated that the effects of pregnancy-induced alveolar differentiation—such as changes in the epithelial architecture of the mammary gland or a general decrease in the hormone responsiveness of the epithelium—may contribute to reduced risk (*2, 8*). While estrogen production by adipose tissue is a major mechanism proposed to contribute to the increased risk of postmenopausal breast cancer in obese women (*13*), far less is known about the underlying mechanisms that link obesity and the decreased risk of ER+/PR+ breast cancer in premenopausal women.

One challenge for understanding the relationship between hormone signaling, pregnancy, and BMI in the healthy human breast is that many of the effects of ovarian hormones within the breast are indirect. The estrogen and progesterone receptors (ER/PR) are expressed in only 10-15% of cells within the epithelium (*14*), and most of the effects of hormone receptor activation are mediated by a complex cascade of paracrine signaling from these hormone-responsive (HR+) luminal cells to other cell types in the breast. Thus, mechanisms for decreased hormone responsiveness in the parous breast could include either: 1) a change in the hormone signaling response of HR+ luminal cells—due to either changes in HR+ luminal cells themselves or non-cell autonomous changes in hormone levels or availability—and/or 2) a reduction in the proportion of HR+ luminal cells, leading to dampened paracrine signaling to other cell types downstream of ER/PR activation. Single-cell analysis tools such as RNA sequencing (scRNAseq) are particularly well-suited for investigating this problem, since they enable unbiased classification of the full repertoire of cell types within the human breast together with their transcriptional state.

Here, we use scRNAseq of twenty-eight premenopausal reduction mammoplasty tissue specimens, together with FACS and immunostaining in an expanded cohort (table S1, N = 44 total samples), to directly measure sample-to-sample variability in cell proportions and cell signaling state in the human breast (Fig. 1A). We develop a computational approach that leverages the inter-sample transcriptional heterogeneity in our dataset to identify coordinated changes in transcriptional states across multiple cell types in the breast. Based on this, we identify a set of correlated gene expression programs in HR+ luminal cells and other cell types representing the paracrine signaling networks activated in response to hormones. Second, we find that prior history of pregnancy is associated with striking changes in epithelial composition, and we propose that these changes are consistent with the protective effect of pregnancy on lifetime breast cancer risk. Finally, we show that paracrine signaling from HR+ luminal cells to neighboring cells depends on both the magnitude of the hormone-signaling transcriptional response and the overall abundance of HR+ cells. Pregnancy and obesity both lead to decreased hormone responsiveness in the breast through these two distinct mechanisms: pregnancy directly affects hormone signaling in HR+ luminal cells whereas obesity reduces the proportion of HR+ luminal cells. Overall, these results provide a comprehensive map of the cycling human breast and identify cellular changes that underlie breast cancer risk factors.

**Fig. 1.**
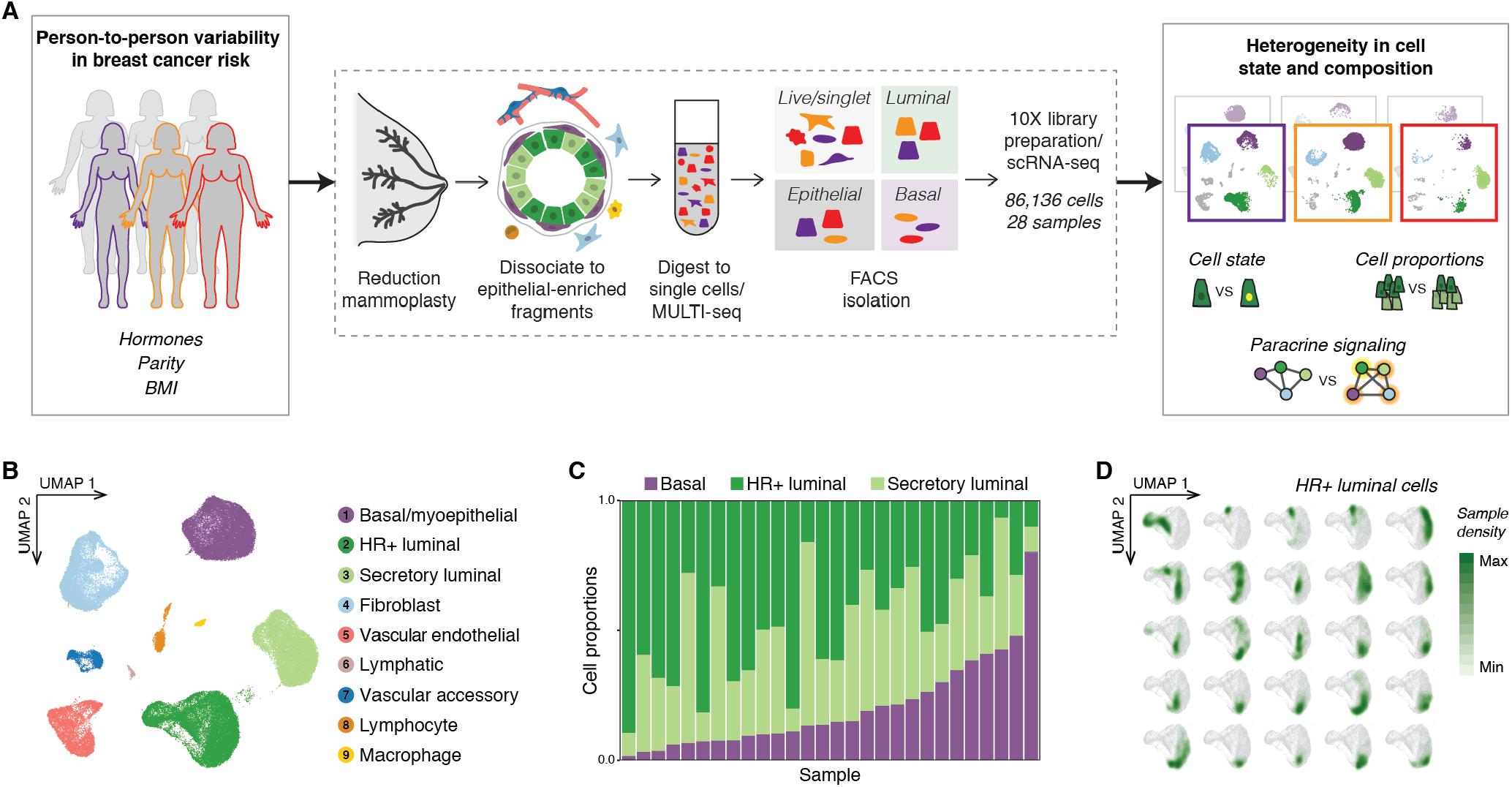
Sample-to-sample variability in epithelial cell proportions and transcriptional cell state in the human breast. (A) Single-cell transcriptional analysis links breast cancer risk factors (parity, BMI, hormones) with person-to-person heterogeneity in cell state, cell proportions, and paracrine signaling. scRNAseq workflow: Reduction mammoplasty samples were processed to epithelial-enriched tissue fragments, followed by processing to single cells and MULTI-seq sample barcoding, FACS isolation, and library preparation. (B) UMAP dimensionality reduction and unsupervised clustering of the combined data from twenty-eight samples identifies the major epithelial and stromal cell types in the breast. (C) Stacked bar plot of the proportion of epithelial cells (HR+ luminal; secretory luminal; basal/myoepithelial) across breast tissue samples. (D) Density plots highlighting the transcriptional cell state of HR+ luminal cells from individuals with at least 100 cells in this cluster.

## Results

### Inter-sample variability in epithelial cell proportions and transcriptional cell state in the human breast

To identify inter-individual differences in cell composition and cell state in the human breast, we performed scRNAseq analysis on 86,136 cells from reduction mammoplasties in 28 premenopausal donors (Fig. 1A and table S1). To obtain an unbiased snapshot of the epithelium and stroma, we collected live/singlet cells from all samples. For a subset of samples, we also collected purified epithelial cells or purified luminal and basal/myoepithelial cells (fig. S1A, table S2). We used MULTI-seq barcoding and *in silico* genotyping for sample multiplexing to minimize technical variability between samples (fig. S1B, *methods*) (*15, 16*).

Sorted basal and luminal cell populations were well-resolved by uniform manifold approximation and projection (UMAP) (fig. S1C). Unsupervised clustering identified one basal/myoepithelial cluster (cluster 1), two luminal clusters (clusters 2-3), and six stromal clusters (clusters 4-9) (Fig. 1B). Based on the expression of known markers, the two luminal clusters were annotated as hormone-responsive (HR+) and secretory luminal cells, and the six stromal clusters were annotated as fibroblasts, vascular endothelial cells, lymphatic endothelial cells (“lymphatic”), smooth muscle cells/pericytes (“vascular accessory”), lymphocytes, and macrophages (Fig. 1B and fig S2A-B). The luminal populations described here closely match those identified as “hormone-responsive/L2” and “secretory/L1” in a previous scRNAseq analysis of the human breast (*17*), as well as microarray data for sorted EpCAM^+^/CD49f^−^ “mature luminal” and EpCAM^+^/CD49f^+^ “luminal progenitor” populations (*18*). Here, we use the nomenclature “hormone-responsive/HR+” and “secretory” to refer to these two cell types. The HR+ cluster was enriched for the hormone receptors ESR1 and PGR (fig. S2C), and other known markers such as ANKRD30A (fig. S2A-B) (*17*). Consistent with previous studies demonstrating variable hormone receptor expression across the menstrual cycle (*19*), expression of ESR1 and PGR transcripts were sporadic and often non-overlapping. Within the HR+ luminal cluster, 22% of the cells had detectable levels of ESR1 or PGR, with only 2% of cells expressing both transcripts (fig. S2D).

Beyond identifying the major cell types, single-cell analysis additionally resolved two sources of inter-sample variability in the human breast. First, while cells from different individuals were represented across all clusters (cluster entropy = 0.93, *methods*) (fig. S3A), the proportions of epithelial cell types were highly variable between samples (Fig. 1C). Across individuals, epithelial cell proportions in the live/singlet and epithelial sort gates ranged from 2-80% for basal/myoepithelial cells, 7-89% for HR+ luminal cells, and 9-70% for secretory luminal cells (fig. S3B). Second, independent of variation in cell proportions, individuals displayed distinct transcriptional signatures within cell types (Fig. 1D and fig S3C). This variation in cell state was not due to technical variability across batches, as cells from the same sample were more similar to each other than cells from different samples, regardless of the day of processing (fig. S3, D and E, table S2, and *methods*).

### Parity is associated with an increased proportion of basal/myoepithelial cells in the epithelium

The breast undergoes a major expansion of the mammary epithelium during pregnancy, followed by a regression back towards the pre-pregnant state after weaning in a process called involution. However, the epithelial architecture remains distinct from that of women without prior pregnancy, consisting of larger terminal ductal lobular units (TDLUs) containing greater numbers of acini. At the same time, individual acini are reduced in size (*2*). We hypothesized that these architectural changes following pregnancy would contribute to differences in epithelial cell proportions between samples.

We focused our initial analysis on the 63,583 cells in the live/singlet and epithelial sort gates to get an unbiased view of how the epithelial composition of the breast changes with pregnancy. The proportion of basal/myoepithelial cells in the epithelium was approximately two-fold higher in women with prior history of pregnancy (parous) relative to women without prior pregnancy (nulliparous) (Fig. 2A and fig. S4A; FDR < 0.02, Wald test). We confirmed these results in an expanded cohort of samples using three additional methods. First, we measured basal cell proportions by flow cytometry analysis of EpCAM and CD49f (fig. S1A). Consistent with clustering results, parity was associated with an increase in the average proportion of EpCAM^−^/CD49f^+^ basal cells from 12% to 39% of the epithelium (Fig. 2B; p < 0.0001, Mann-Whitney test). The proportion of basal cells did not vary with other discriminating factors such as BMI, race, or hormonal contraceptive use, but was weakly associated with age (R^2^ = 0.20, p < 0.04, Wald test) (fig. S4B). To determine the relative effect of each factor, we performed multiple linear regression analysis and found that the basal cell fraction positively correlated with pregnancy history (p < 2e-05, Wald test), but not age (p = 0.17, Wald test) (Table S3; R^2^ = 0.77, p < 8e-6). Next, as FACS processing steps may affect tissue composition, we performed two further analyses. We reanalyzed two previously published microarray datasets of total RNA isolated from core needle biopsies from either premenopausal (n = 71 parous/42 nulliparous) or postmenopausal (n = 79 parous/30 nulliparous) women (*20, 21*), and confirmed a significant increase in the basal/myoepithelial markers KRT5, KRT14, and TP63 relative to luminal markers in parous samples (fig. S4C). Finally, we performed immunostaining and confirmed an approximately 2-fold increase in the ratio of p63+ basal cells to KRT7+ luminal cells in intact tissue sections (Fig. 2C; p < 0.001, Mann-Whitney test). Notably, immunostaining demonstrated that this change in epithelial proportions was specific to TDLUs rather than ducts (fig. S5A). We hypothesized that the increased frequency of basal/myoepithelial cells observed in parous women could be explained, in part, by changes in TDLU architecture. To test this, we performed a morphometric comparison of TDLUs between parous and nulliparous samples in our dataset. Consistent with previous reports (*2*), we observed a marked decrease in the average diameter of individual acini in parous women (fig. S5B; p < 0.002, Mann-Whitney test). Additionally, we found that the average thickness of the luminal cell layer was linearly associated with acinus diameter (fig. S5C; R^2^ = 0.89, p < 0.0001) and reduced in parous women (fig. S5D; p < 0.002, Mann-Whitney test).

**Fig. 2.**
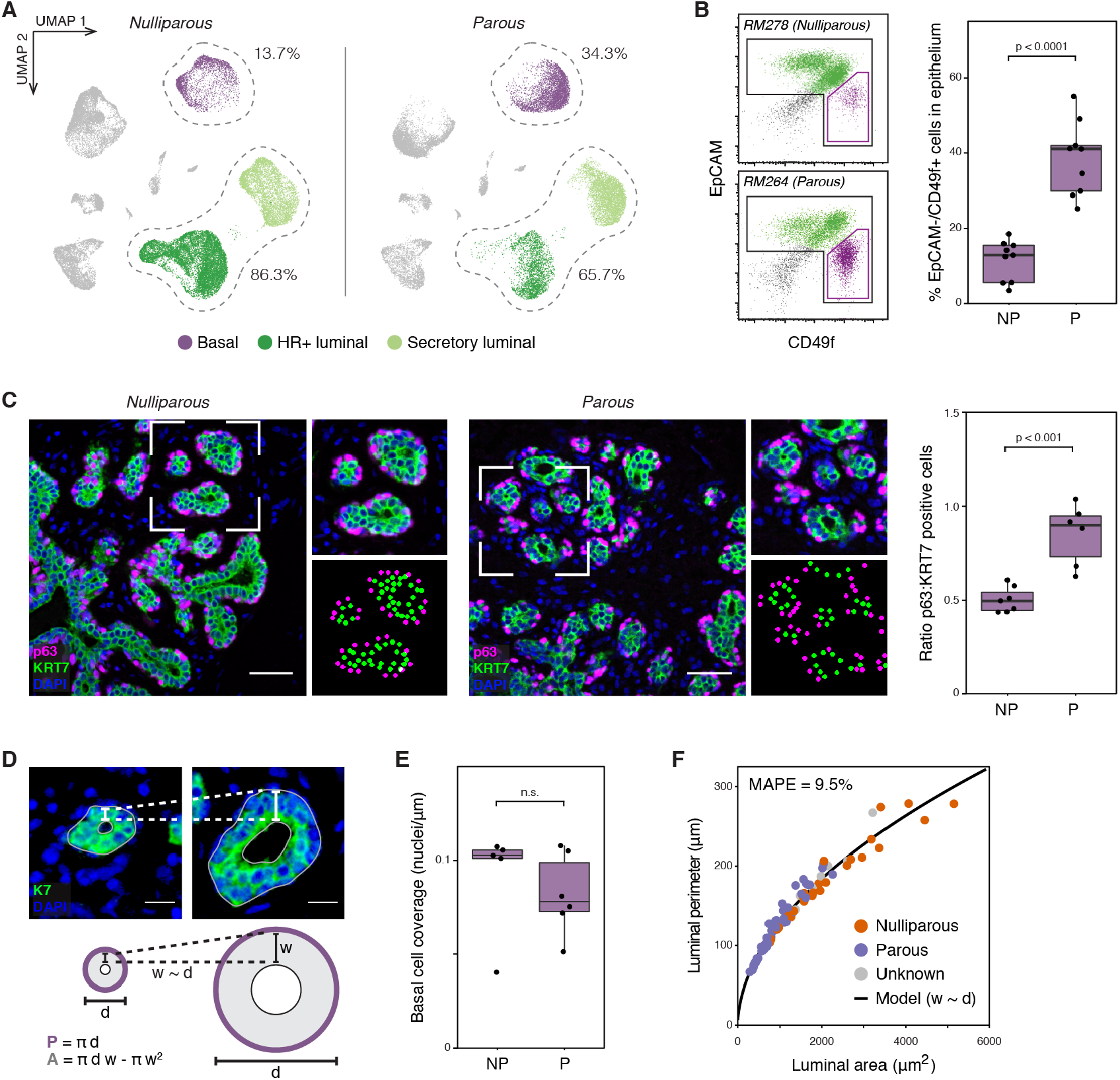
Prior history of pregnancy is associated with an increased proportion of basal cells in the mammary epithelium. (A) UMAP plot of sorted live singlet and epithelial cells from nulliparous and parous samples, with the percent of luminal and basal/myoepithelial cells highlighted. (B) Representative FACS analysis of the percentage of EpCAM^−^/CD49f^+^ basal cells within the Lin^−^ epithelial population, and quantification of the percentage of basal cells in parous (P) versus nulliparous (NP) women (n = 18 samples; p < 0.0001, Mann-Whitney test). (C) Immunostaining for the basal/myoepithelial marker p63 and pan-luminal marker KRT7, and quantification of the ratio of p63+ basal/myoepithelial cells to KRT7+ luminal cells for samples with or without prior history of pregnancy (NP = nulliparous, P = parous; n = 13 samples; p < 0.001, Mann-Whitney test). Scale bars 50 µm. (D) Two-dimensional geometric model of the relative space available for basal cells (outer perimeter of the luminal layer, P) and luminal cells (area of the luminal layer, A) within individual acini. Acini were modeled as hollow circles with a shell thickness proportional to their diameter. Scale bars 15 µm. (E) Quantification of the average basal cell coverage (nuclei per μm of luminal perimeter) in acini from terminal ductal lobular units (TDLUs) in nulliparous (NP) versus parous (P) samples (p = 0.66, Mann-Whitney test). (F) Results of geometric modeling depicting the relative area and perimeter of the luminal layer as a function of acinus diameter. Dots represent measurements of individual acini from TDLUs in parous (n=53 acini from 7 samples) or nulliparous (n=29 acini from 7 samples) women as indicated (mean absolute percentage error = 9.5%).

To determine how these parameters influence the relative proportions of each cell type, we implemented a simple geometric model (Fig. 2D, *methods*). Surprisingly, when normalized to cross-sectional area (for luminal cells) or perimeter (for basal cells), there was no change in luminal cell density or basal cell coverage between parous versus nulliparous samples (Fig. 2E and fig. S5E). Across all samples, the number of basal or luminal cells per acinus was proportional to the space available for each cell type (fig. S5F; p < 0.0001, Wald test). However, geometric modeling accurately predicted the relationship between the luminal area and outer perimeter for individual acini (mean absolute percentage error loss = 9.5%) and demonstrated that as individual acini increased in size, the space available for luminal cells (luminal area) increased at a faster rate than the space available for basal cells (luminal perimeter) (Fig. 2F). Thus, the observed differences in epithelial cell proportions between parous and nulliparous samples are not due to a change in basal/myoepithelial coverage, but rather a change in the overall morphology of the luminal layer and relative surface area of individual acini in parous women.

### Obesity is associated with a reduction in the proportion of HR+ luminal cells

While parity was associated with a decreased overall proportion of luminal cells in the epithelium, the proportions of individual HR+ and secretory subtypes within the luminal compartment were highly variable. Consistent with previous work (*5, 22*), we observed reduced frequencies of HR+ luminal cells in parous women (fig. S4A; FDR < 0.03, Wald test with post hoc multiple-comparisons test). However, the proportion of secretory luminal cells was not associated with parity (fig. S4A). Together, these data suggested that additional factors influence the relative proportion of HR+ versus secretory cells within the luminal compartment. We therefore performed multiple comparison analysis to test for the effects of parity, BMI, race, age, and hormonal contraceptive use on the proportions of HR+ versus secretory luminal cells. We found that the relative proportion of HR+ luminal cells was reduced in obese women (BMI ≥ 30) (Fig. 3A; FDR < 0.0002, Wald test with post hoc multiple-comparisons test) and did not vary significantly with other discriminating factors such as age, reproductive history, hormonal contraceptive use, or race (fig. S6A; Wald test with post hoc multiple-comparisons test). On a continuous scale, each 12 units of BMI was associated with a 2-fold reduction in the proportion of HR+ cells in the luminal compartment (fig. S6B; FDR < 0.001, Wald test with post hoc multiple-comparisons test). We observed similar results using clustering analysis from the 10,795 cells in the luminal sort gate (fig. S6C).

**Fig. 3.**
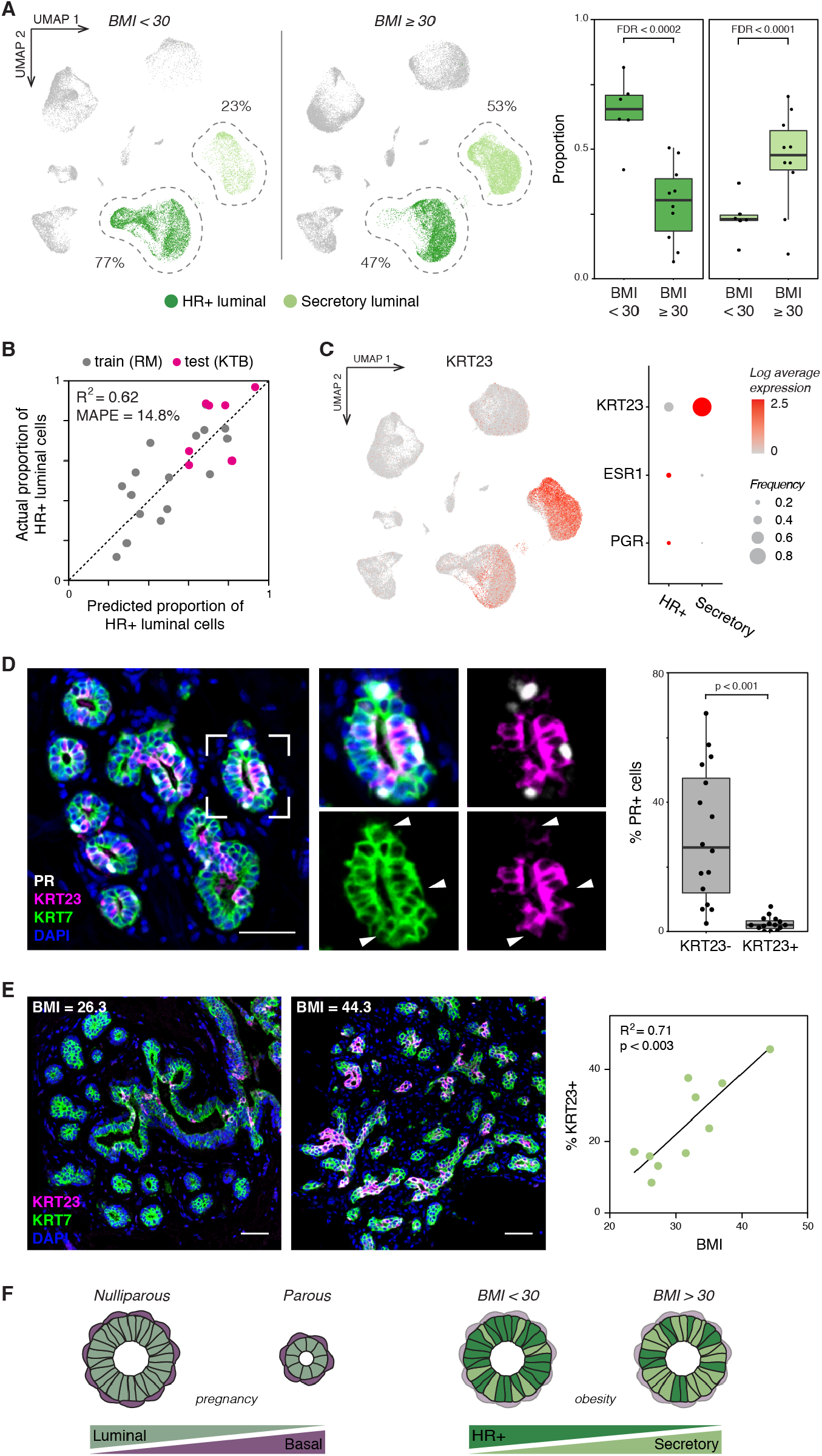
Obesity is associated with a decreased proportion of hormone-responsive cells in the luminal compartment. (A) *Left:* UMAP plot of sorted live/singlet and epithelial cells from non-obese (BMI < 30) and obese (BMI ≥ 30) samples, highlighting hormone-responsive (HR+) and secretory luminal cells. *Right:* Quantification of the proportion of HR+ or secretory cells in the luminal compartment of obese versus non-obese samples (n = 16 samples; FDR < 0.0002, Wald test). (B) A quasi-Poisson regression model accurately predicts the proportion of HR+ cells in the luminal compartment as a function of BMI in an independent cohort of Komen Tissue Bank core biopsy samples (mean absolute percentage error = 14.8%; see also fig. S6B). (C) *Left:* UMAP depicting log normalized expression of KRT23. *Right:* Dot plot depicting the log normalized average and frequency of KRT23, ESR1, and PGR expression across luminal cell types. (D) Co-immunostaining of PR, KRT23, and the pan-luminal marker KRT7, and quantification of the percentage of PR+ cells within the KRT23- and KRT23+ luminal cell populations (n = 16 samples; p < 0.001, Mann-Whitney test). (E) Co-immunostaining of KRT23 and KRT7 and linear regression analysis of the percentage of KRT23+ luminal cells versus BMI (n = 10 samples; R^2^ =0.71, p < 0.003, Wald test). Scale bars 50 µm. (F) Summary of changes in epithelial cell proportions with pregnancy and obesity.

One limitation of the reduction mammoplasty dataset was that all samples classified as non-obese were from nulliparous women less than 24 years old, whereas obese samples were more likely to be from parous and older age women (table S1, fig S7A). Therefore, we performed scRNAseq analysis on an independent set of breast core biopsies from healthy premenopausal women who donated tissue to the Komen Tissue Bank (KTB) (fig. S7, B-E; table S2). In contrast with the reduction mammoplasty cohort, the KTB cohort consisted of older (37-47 years) parous samples with BMI in the normal or overweight range (BMI 20.7-28.3) (table S1, fig. S7A). Using the reduction mammoplasty cohort as a training set, we accurately predicted the proportion of HR+ luminal cells in the KTB cohort as a function of BMI with a mean absolute percentage error of 14.8% (Fig. 3B).

To verify these results in tissue sections, we performed immunostaining for ER and PR. There was a trend toward decreased expression of PR with increasing BMI, but the change was not statistically significant (p = 0.11, Wald test; fig. S8A). Notably, ER and PR protein expression was variable and partly non-overlapping, ranging from 11-71% overlap (fig. S8B). As we had previously also observed heterogeneous expression of ESR1 and PGR transcripts within the HR+ luminal cell cluster (fig. S2, C and D), we hypothesized that the variability in staining was due to changes in ER/PR expression, stability, and nuclear localization that have all been previously observed based on hormone receptor activation status (*19, 23, 24*). Based on this, we predicted that ER/PR transcript and protein levels would co-vary across samples due to the overall proportion of HR+ luminal cells and their hormonal microenvironment, but would be stochastically expressed in individual cells at any one time due to fluctuations in mRNA and protein expression, localization, and stability. To test this, we performed co-immunostaining and RNA-FISH and confirmed that although ER transcript and protein levels correlate across tissue sections (R^2^ = 0.60, p < 0.01), they do not correlate on a per-cell basis (p = 0.63, Wilcoxon signed-rank test)—on average, only 31% of cells expressing ESR1 transcript also expressed ER protein (fig. S8C). Importantly, expression of ESR1 or PGR transcript was highly specific for cells in the HR+ luminal cluster, although the sensitivity of each transcript for the HR+ cluster was low and varied across individuals (fig. S8D). Thus, these data demonstrate that immunostaining for nuclear hormone receptors underestimates the fraction of cells in the HR+ lineage and that lack of ER/PR expression cannot be used to reliably define a cell as part of the secretory versus HR+ luminal cell lineages.

On the basis of these results, we sought to identify another marker to distinguish between luminal subpopulations, and identified keratin 23 (KRT23) as highly enriched in the secretory luminal cell cluster (Fig. 3C), as was also reported by a previous scRNAseq study (*17*). Immunohistochemistry for KRT23 and PR or ER confirmed that these proteins are expressed in mutually exclusive luminal populations (Fig. 3D, and fig. S9, A and B; p < 0.001 and p < 0.01, Mann-Whitney test). The proportion of KRT23+ luminal cells in each sample was also highly correlated with the proportion of secretory luminal cells identified by scRNAseq (fig. S9C; R^2^ = 0.71, p < 0.0001, Wald test). KRT23 thus represents a discriminatory marker between the two luminal populations. Staining in intact tissue sections confirmed that the proportion of KRT23+ secretory luminal cells increased by about 17% for every 10-unit increase in BMI (Fig. 3E; R^2^ = 0.71, p < 0.003). Together, these data demonstrate that there are two independent effects of reproductive history and body weight on cell proportions in the mammary epithelium: parity affects the ratio of basal to luminal cells whereas BMI affects the ratio of HR+ versus secretory luminal cells (Fig. 3F).

### Hormone signaling is a primary axis of transcriptional variability in HR+ luminal cells

Beyond differences in cell proportions, we found that transcriptional cell state within clusters was a second source of inter-sample variability in our dataset (Fig. 1D, and fig. S3, C and E). Since estrogen and progesterone are master regulators of breast development, and the levels of these hormones fluctuate across the menstrual cycle, we hypothesized that hormone signaling would represent a major source of transcriptional heterogeneity across samples. Consistent with this, we previously observed a high degree of sample-to-sample variation in ER/PR expression (fig. S8D) within the HR+ luminal cell cluster, which has been shown to vary based on hormone receptor activation state (*19, 23, 24*).

To quantify cell state in HR+ luminal cells, we first performed principal component (PC) analysis on this population. Analysis of ranked gene loadings demonstrated that variation across PC1 in HR+ cells was driven by genes involved in the response to hormone receptor activation, including the essential PR target genes WNT4 and TNFSF11 (RANKL) (*6, 25*) (Fig. 4A). Of the 20 genes with the highest loadings in PC1, 12 have been previously described as associated with either estrogen signaling, progesterone signaling, or the luteal phase of the menstrual cycle when progesterone is at its peak (fig. S10A, table S4) (*6, 26*-*35*). Thus, transcriptional changes associated with hormone signaling state (PC1) are a dominant source of variation in HR+ luminal cells (fig. S10B).

**Fig. 4.**
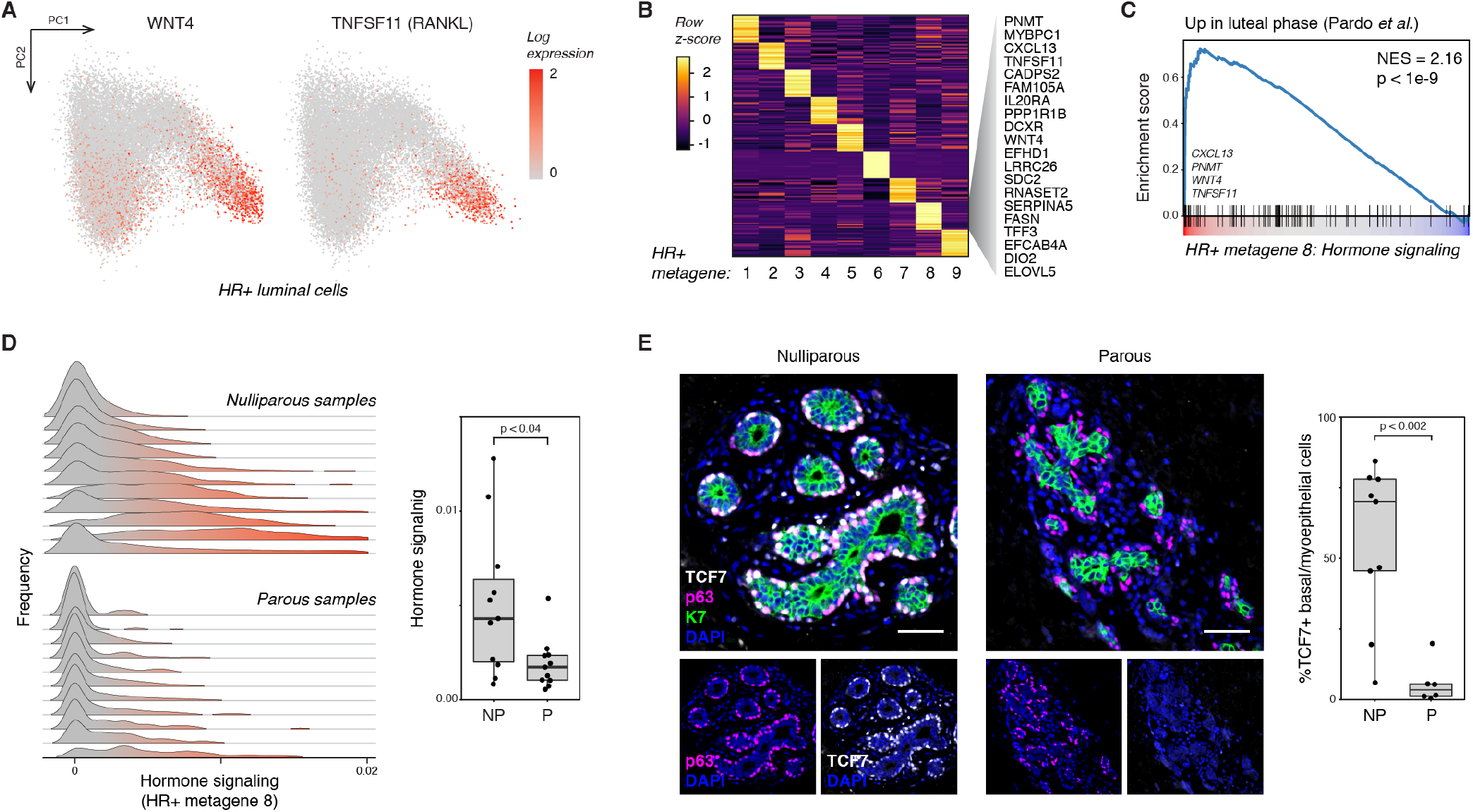
Hormone signaling is a primary axis of transcriptional variability in HR+ luminal cells. (A) PCA plot of HR+ luminal cells depicting expression of WNT4 and TNFSF11 (RANKL) in log normalized counts. (B) Non-negative matrix factorization identifies a specific gene signature of hormone signaling in HR+ luminal cells. Heatmap depicting the top 20 genes expressed in each HR+ cell metagene, highlighting marker genes in HR+ metagene 8. (C) Gene set enrichment analysis of HR+ cell metagene 8, showing enrichment of genes shown to be upregulated during the luteal phase of the menstrual cycle (NES = 2.16, p < 1e-9) (*29*). (D) Ridge plots depicting the distribution of HR+ metagene 8 (hormone signaling) expression in HR+ luminal cells across samples, and quantification of the average expression of metagene 8 in nulliparous (NP) versus parous (P) samples (n = 22 samples, p = 0.04, Mann-Whitney test). (E) Immunostaining for TCF7, p63, and KRT7, and quantification of the percentage of TCF7+ cells within the p63+ basal/myoepithelial cell compartment for nulliparous (NP) versus parous (P) samples (n=15 samples; p < 0.002, Mann-Whitney test).

As PC analysis seeks to maximize the variance of a projected dataset, it may combine gene signatures from multiple transcriptional states into a single component (fig. S10A) (*36*). Therefore, we performed non-negative matrix factorization (NMF) to identify a specific gene signature of hormone signaling, and identified 9 distinct gene expression programs, or “metagenes” in HR+ luminal cells (Fig. 4B, and fig. S10, C and D) (*37, 38*). Cell embedding in PC1 was highly correlated with expression of metagene 8 (Pearson correlation = 0.79, fig. S10E). Analysis of ranked gene loadings demonstrated that this “hormone signaling” metagene comprised a similar gene expression program as PC1, including the PR targets WNT4 and TNFSF11 (RANKL) and the ER target TFF3 (Fig. 4B). The hormone signaling metagene was highly enriched for genes upregulated during the luteal phase of the menstrual cycle (Fig. 4C; p < 1e-9) (*29*), and for transcripts in the Molecular Signatures Database Hallmark “early estrogen response” (p < 0.006) and “late estrogen response” (p < 0.007) gene sets (fig. S10F) (*39*). Thus, NMF identified a distinct transcriptional signature for hormone receptor activation in HR+ luminal cells, representing both known hormone-responsive genes as well as new markers of the hormone response specifically in the HR+ luminal cell population.

### The hormone signaling response of HR+ luminal cells is reduced in parous women

Previous epidemiologic analyses have demonstrated that the protective effect of parity against breast cancer is specific for ER+/PR+ tumors (*12*). Decreased hormone responsiveness following pregnancy is one proposed mechanism for this effect (*8*). Supporting this, previous studies demonstrated decreased expression of the PR effector WNT4 following pregnancy (*5, 22*). Moreover, in an explant culture model, estrogen consistently induced expression of the ER target gene AREG only in nulliparous women (*4*). As the effects of hormones in the breast are primarily mediated by paracrine signaling from HR+ luminal cells, this decreased hormone responsiveness could be caused by either: 1) a change in the magnitude of paracrine signals produced by each HR+ luminal cell, and/or 2) a reduction in the overall proportion of HR+ luminal cells leading to a “dilution” of paracrine signals following ER/PR activation. It has been difficult to distinguish between these mechanisms using bulk tissue-level analyses. By probing the single-cell transcriptional landscape of the HR+ luminal cell population, NMF analysis provided a means to directly interrogate whether parity influences the per-cell hormone signaling response of HR+ luminal cells.

To quantify variation in hormone signaling in HR+ luminal cells, we first measured the similarity between each sample’s single-cell distribution across metagene 8. Hierarchical clustering identified two sets of samples, representing high or low hormone signaling (fig. S11A). Based on this, we found that while the level of hormone signaling in HR+ luminal cells varied between nulliparous women, likely reflecting differences in hormone levels across the menstrual cycle or due to hormonal contraceptive use, per-cell hormone signaling in HR+ luminal cells was significantly reduced in parous women (p < 0.04, Mann-Whithney test; Fig. 4D, and fig. S11B). Importantly, equal numbers of individuals from each cohort were using hormonal birth control (n = 4 out of 11 nulliparous or parous individuals, table S1). For women not using hormonal birth control (n = 7 out of 11 nulliparous or parous individuals), we modeled the expected number of samples with high hormone signaling based on a binomial distribution using average menstrual cycle phase lengths (*40*). The number of nulliparous samples with high hormone signaling matched the expected modal number of samples in the luteal phase (3 of 7 samples, P = 0.29), whereas the number of parous samples with high hormone signaling was significantly lower than expected based on the average length of the follicular and luteal phases of the menstrual cycle (0 of 7 samples, P = 0.02) (fig. S11C). Thus, the decreased per-cell hormone signaling seen in HR+ luminal cells from parous women cannot be explained by differences in hormonal contraceptive use or random sampling across the menstrual cycle.

To identify differentially expressed genes between nulliparous and parous women with high sensitivity, we generated a pseudo-bulk dataset of aggregated HR+ luminal cells from each sample (*methods*) and confirmed that parous women had decreased expression of the canonical hormone-responsive genes AREG, WNT4, PGR, TNFSF11 (RANKL), and TFF1 (fig. S11D, table S5). Notably, the progesterone receptor itself is an ER target gene (*41*). Staining for the progesterone receptor and K23 confirmed that PR expression was reduced in the HR+ luminal cell subpopulation (K7+/K23-) of parous samples (fig. S11E). Finally, we confirmed that paracrine signaling downstream of PR activation was specifically reduced in parous samples by assessing the effects of one of these genes, WNT4. As WNT4 from HR+ luminal cells has been shown to signal to basal cells (*25*), we performed co-immunostaining for the WNT effector TCF7 and basal/myoepithelial cell marker p63 and found that TCF7 expression was markedly decreased in parous samples (p < 0.002, Mann-Whitney test; Fig. 4E). This decrease was not due to differences in epithelial architecture, as TCF7 staining in ducts versus TDLUs within the same samples was unchanged (fig. S11F). Together, these data demonstrate that transcriptional variation among HR+ luminal cells is primarily related to hormone signaling, that transcription along this axis (HR+ metagene 8) is reduced in women with prior history of pregnancy, and that these transcriptional changes coincide with a reduction in downstream paracrine signaling to basal/myoepithelial cells.

### Identification of coordinated changes in signaling states across cell types in the breast downstream of hormone signaling

As the effects of estrogen and progesterone on other cell types in the breast are controlled by paracrine signaling from HR+ luminal cells, we reasoned that hormone receptor activation in HR+ luminal cells would be correlated with transcriptional changes in other cell types, representing the downstream paracrine response. To identify putative transcriptional signatures of the paracrine response, we developed a computational framework that leverages the person-to-person transcriptional heterogeneity observed within cell types to find coordinated changes in cell signaling states across samples. This approach builds upon previous studies that used heterogeneous expression of individual genes across tissue regions (*42*) or small cell populations (*43, 44*) to identify co-regulated transcripts. However, rather than identifying individual genes that co-vary either spatially or within cell populations, we instead identify distinct transcriptional cell states (metagenes) within cell types that co-vary across samples. First, we decomposed each cell type into sets of gene expression programs, or “metagenes”, using NMF as described above (fig. S10, C and D, and fig. S12, A and B). We then quantified the average expression of each cell type-specific metagene for each sample and constructed a weighted network of coordinated gene expression programs based on the pair-wise Pearson correlations between metagenes (fig. S12C, *methods*). Finally, we identified modules of highly correlated gene expression programs using the infomap community detection algorithm (*45*). Using this approach, we identified three major modules— annotated here as “resting state”, “paracrine signaling”, and “involution” modules—comprising highly interconnected transcriptional states across cell types in the breast (Fig. 5A).

**Fig. 5.**
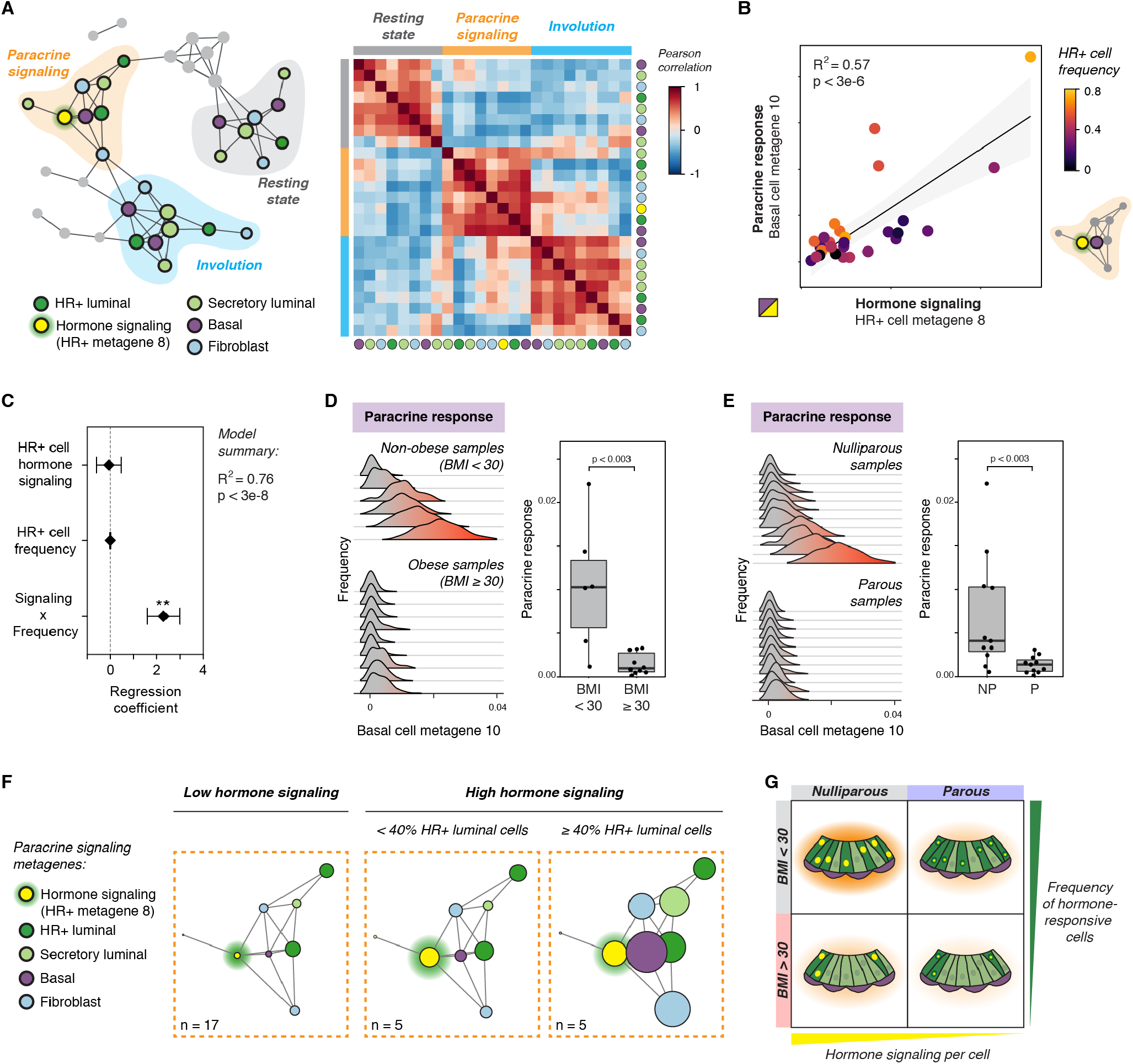
Identification of coordinated changes in signaling states across cell types in the breast. (A) *Left:* Network graph of correlated gene expression programs in the human breast. Nodes represent distinct metagenes in the indicated cell types, and edges connect highly correlated metagenes (Pearson correlation coefficient > 0.5, p < 0.05). Modules of correlated gene expression programs were identified using the infomap community detection algorithm. The “hormone signaling” metagene in HR+ cells (HR+ metagene 8) is highlighted in yellow. *Right:* Heatmap depicting Pearson correlation coefficients between metagenes in the three major modules. (B) Linear regression analysis of basal cell state across metagene 10 (paracrine response) versus HR+ luminal cell state across metagene 8 (hormone signaling) (R^2^ = 0.57, p < 3e-6, Wald test). Dots represent the average expression of each metagene within a sample, colored by the proportion of HR+ luminal cells in the epithelium for that sample. (C) Summary of multiple linear regression analysis with three predictors: HR+ cell hormone signaling (HR+ metagene 8), the frequency of HR+ cells in the epithelium, and an interaction term representing the combined effects of HR+ signaling and frequency (Signaling × Frequency). (D) Ridge plots depicting the distribution of basal cell metagene 10 (paracrine response) expression across samples, and quantification of the average expression in obese (BMI ≥ 30) versus non-obese (BMI < 30) samples (n = 16 samples, p < 0.003, Mann-Whitney test). (E) Ridge plots depicting the distribution of basal cell metagene 10 (paracrine response) expression across samples, and quantification of the average expression in nulliparous (NP) versus parous (P) samples (n = 22 samples, p < 0.003, Mann-Whitney test). (F) Network subgraph of the “paracrine signaling” module with node sizes proportional to the mean expression of each metagene in: samples with low hormone signaling (n = 17, see fig. S11A), samples with high hormone signaling and < 40% HR+ luminal cells in the epithelium (n = 5), or samples with high hormone signaling and ≥ 40% HR+ luminal cells in the epithelium (n = 5). Nodes represent distinct metagenes in the indicated cell types, and edges connect highly correlated metagenes (Pearson correlation coefficient > 0.5; p < 0.05). (G) Schematic depicting how parity and obesity lead to decreased hormone signaling in the breast through distinct mechanisms. Parity directly affects the per-cell hormone response in HR+ luminal cells, whereas BMI leads to a reduction in the proportion of HR+ luminal cells in the epithelium.

The “resting state” module consisted of gene expression programs that were negatively correlated with hormone signaling in HR+ luminal cells (Fig. 5A, and fig. S13A). Metagenes in this module were primarily enriched for pathways involved in ribosome biogenesis and RNA processing (fig. S14B). The “paracrine signaling” module comprised gene expression programs that were positively correlated with hormone signaling in HR+ luminal cells (Fig. 5A, and fig. S14A). As expected based on the central role HR+ cells play in the response to estrogen and progesterone, the HR+ hormone signaling metagene (HR+ metagene 8) had the greatest influence on information flow within this module, as measured by betweenness centrality (fig. S14A). Our analysis revealed that high levels of hormone signaling in HR+ cells coincided with the emergence of a second transcriptional state—HR+ metagene 5—in a distinct subpopulation of HR+ luminal cells (fig. S14B). Marker analysis and gene set enrichment analysis demonstrated that HR+ metagene 5 was characterized by upregulation of a hypoxia gene signature and pro-angiogenic factors such as VEGFA and ANGPTL4 (fig. S14C). The identification of this “hypoxia” gene signature is consistent with a previous study using microdialysis of healthy human breast tissue, which found that VEGF levels increased in the luteal phase of the menstrual cycle (*46*). As estrogen response elements have been identified in the untranslated regions of VEGFA (*47*), our results suggest that this increased expression may be, in part, a direct effect of hormone signaling to this subpopulation of HR+ cells.

We next investigated gene expression programs in other epithelial and stromal populations that were highly correlated with hormone signaling in HR+ luminal cells. NMF and network analysis identified a subpopulation of proliferative secretory luminal cells within the paracrine signaling module (fig. S14D). This “proliferation” metagene was highly enriched for cell-cycle related genes (fig. S14D) previously found to be upregulated during the luteal phase of the menstrual cycle (fig. S14E) (*29*). Moreover, similar to HR+ cells, basal/myoepithelial cells in samples with high levels of hormone signaling had enrichment of transcripts involved in hypoxia and angiogenesis such as VEGFA and ANGPTL4 (fig. S14F). Gene set enrichment analysis demonstrated that variation across this basal cell “paracrine response” metagene was driven by genes involved in epithelial-mesenchymal transition, cell adhesion, cell motility, and extracellular matrix (ECM) organization (fig. S14F), suggesting that changes in basal/myoepithelial cell motility, cell-cell, and cell-ECM interactions underlie the previously reported morphological changes observed in the breast epithelium across the menstrual cycle (*48*). We observed similar morphological changes by H&E staining in samples classified as having high versus low hormone signaling, including the emergence of more distinct luminal and myoepithelial cell layers (fig S14G). Finally, previous studies have identified alterations in stromal organization and ECM composition across the menstrual cycle (*49, 50*). Consistent with this, hormone signaling in HR+ luminal cells correlated with two distinct gene expression programs in fibroblasts: a “tissue remodeling” metagene characterized by upregulation of ECM proteins including collagens (COL3A1, COL1A1, COL1A2) and fibronectin (FN1), and a “proinflammatory” gene expression program representing upregulation of cytokines and growth factors such as IL6 and TGFB3 (fig. S14H).

Finally, gene set enrichment analysis of the third module uncovered a transcriptional signature in HR+ and secretory luminal cells that was similar to that identified during post-lactational involution (fig. S15, A and B) (*51, 52*). These “involution” metagenes were characterized by high expression of death receptor ligands such as TNFSF10 (TRAIL) and genes involved in the defense and immune response, including interferon-response genes (fig. S15, B and C). The involution signature in secretory luminal cells was also characterized by expression of major histocompatibility complex class II (MHCII) molecules and the phagocytic receptors CD14 and MARCO (fig. S15B), suggesting that these cells play a role as non-professional phagocytes in the clearance of apoptotic cells, similar to what has been described during post-lactational involution (*53*). Previous data have demonstrated that the fraction of apoptotic cells in the mammary epithelium peaks between the late luteal and early follicular phases of the menstrual cycle (*54*). Notably, TGFB3 signaling is a major signaling molecule involved in post-lactational involution that enhances phagocytosis by mammary epithelial cells (*55*), suggesting that TGFB3 secreted by fibroblasts at the end of the luteal phase (fig. S14H) activates a subset of secretory luminal cells during the late luteal/early follicular phase that go on to express “involution” markers including phagocytic receptors.

Together, these results demonstrate how the underlying sample-to-sample variability in scRNAseq data can be used to infer functional connections between cell types in paracrine signaling networks. Using this computational framework, we find that variation in hormone signaling in HR+ luminal cells is linked to transcriptional variability across all major cell types in the breast. Strikingly, many of these changes closely mimic those seen during the pregnancy/involution cycle that have been linked to a transient increased breast cancer risk following pregnancy (*56, 57*).

### The proportion of HR+ luminal cells predicts basal cell paracrine signaling state

Previously, we demonstrated that parity was associated with a change in the per-cell hormone signaling response of HR+ luminal cells (Fig. 4D), whereas increased BMI was associated with a reduction in the proportion of HR+ cells in the luminal compartment (Fig. 3). As the effects of ER/PR activation are controlled by paracrine signaling from HR+ luminal cells to other cell types, we reasoned that the overall proportion of HR+ luminal cells in the epithelium was a second mechanism that could affect the hormone responsiveness of the breast. While the downstream effects of hormone receptor activation in HR+ luminal cells are controlled by a complex set of signaling networks, previous work has shown that HR+ cells signal directly to basal cells via WNT (*25*). Since WNT proteins generally form short-range signaling gradients (*58*), we predicted that the paracrine response in basal cells would be particularly sensitive to reductions in the proportion of HR+ luminal cells. Consistent with this idea, while the basal cell “paracrine response” metagene was linearly associated with the hormone signaling state of HR+ luminal cells (R^2^ = 0.57, p < 3e-6, Wald test), positive outliers tended to have a greater proportion of HR+ luminal cells and negative outliers tended to have a lower proportion of HR+ luminal cells in the epithelium (Fig. 5B).

To formally test this prediction, we modeled the basal cell paracrine response as a linear response to three variables: HR+ cell hormone signaling (HR+ metagene 8), the frequency of HR+ cells in the epithelium, and an interaction term representing the combined effects of HR+ signaling and frequency (Signaling × Frequency). This combined model accounted for over 75% of the sample-to-sample variation across the paracrine response metagene in basal cells (R^2^ = 0.76, p < 3e-8; Fig. 5C, fig. S16A, and table S6). Importantly, only the interaction term (Signaling × Frequency) was a significant predictor of basal cell transcriptional state (p < 0.002, Wald test; Fig. 5C and table S6), demonstrating that the basal cell paracrine response requires both hormone signaling in HR+ cells and an appreciable abundance of HR+ cells in the epithelium. Together, these results are consistent with a model in which the proportion of HR+ luminal cells in the epithelium influences the magnitude of paracrine signaling to basal/myoepithelial cells downstream of estrogen and progesterone.

Based on these results, we predicted that BMI would influence paracrine signaling from HR+ luminal cells to basal/myoepithelial cells, since HR+ luminal cells are reduced in obese women (Fig. 3). Confirming this, we found that while direct hormone signaling in HR+ cells was not significantly affected by obesity (p = 0.31, Mann-Whitney test; fig. S16B), the downstream basal cell paracrine response was significantly reduced in obese samples (p < 0.003, Mann-Whitney test; Fig. 5D). Consistent with the reduced hormone signaling previously observed in HR+ cells from parous women (Fig. S4D), parity was also associated with a reduction in the basal cell paracrine response (p < 0.003, Mann-Whitney test; Fig. 5E).

Gene set enrichment analysis demonstrated that variation in the basal cell “paracrine signaling” metagene was driven by genes involved in contractility and cell motility (fig. S14F). To determine whether these genes were differentially expressed in obese and/or parous women, we generated a “pseudo-bulk” dataset of basal cells from each sample. Of the 195 genes significantly downregulated in parous samples and 148 genes significantly downregulated in obese samples, 68 were reduced across both groups (fig. S16C and table S7). Both parous and obese samples had decreased expression of contractility-related genes including ACTA2, ACTG2, CNN1, MYH11, MYL9, and MYLK, as well as the basement membrane proteins COL4A1 and COL14A1 (fig. S16C and table S7). Finally, consistent with the idea that parity and obesity reduce the paracrine response of basal cells to hormone signaling, expression of the WNT target genes SPP1 and WLS were also reduced in both subsets. Overall, these results support a model in which paracrine signaling downstream of hormone receptor activation depends on both the magnitude of signaling from HR+ luminal cells and their overall abundance (Fig. 5F). Moreover, parity and BMI affect the hormone responsiveness of the breast through these two distinct mechanisms: parity directly alters the per-cell hormone signaling response in HR+ luminal cells, whereas BMI indirectly affects hormone signaling by reducing the proportion of HR+ luminal cells in the mammary epithelium (Fig. 5G). Importantly, both mechanisms have a common effect on the breast microenvironment by determining the relative intensity of the paracrine response in neighboring cell types (Fig. 5, D-F).

## Discussion

In this study, we combine single-cell analyses, immunostaining, and computational modeling to deconstruct the major sources of sample-to-sample heterogeneity in the human breast. We identify inter-sample variation in epithelial cell proportions, hormone sensitivity, and transcriptional state across the breast microenvironment, and link these patterns of heterogeneity to biological variables known to modulate breast cancer risk. By using single-cell measurements, we separate out the effects of variation in cell proportions from variation in transcriptional state. We then developed a computational pipeline that leverages the inter-sample transcriptional heterogeneity in our dataset to identify coordinated changes in cell signaling states across cell types. Using this approach, we identify a set of highly correlated gene expression programs representing the *in situ* response to hormone receptor activation in HR+ cells and the effects of downstream paracrine signaling in other cell types. Furthermore, we show that person-to-person heterogeneity in hormone responsiveness in the breast is directly linked to two factors known to modulate premenopausal breast cancer risk—reproductive history and BMI.

Pregnancy has a pronounced protective effect against breast cancer, with up to a 50% reduction in breast cancer risk for women with multiple full-term pregnancies at a young age (*8*). Our analysis revealed that parity is associated with a stark increase in the proportion of basal and/or myoepithelial cells within the breast epithelium. Previous work has described two tumor-protective features of myoepithelial cells: they are resistant to malignant transformation (*59, 60*) and also act as a natural and dynamic barrier that prevents tumor cell invasion (*61, 62*). Thus, our data suggest that pregnancy protects against breast cancer risk both by decreasing the relative frequency of luminal cells—the tumor cell-of-origin for most breast cancer subtypes (*63*-*65*)—and by suppressing progression to invasive carcinoma.

Lifetime hormone exposure is another major determinant of breast cancer risk (*1*). Here, we use matrix decomposition and network analysis to map the coordinated changes in cell state that occur in response to paracrine signaling from HR+ luminal cells. Strikingly, many of these changes closely mimic those seen during the pregnancy/involution cycle that have been linked to a transient increased breast cancer risk following pregnancy (*56, 57, 66*). First, we identify a proliferative gene signature in secretory luminal cells that is highly correlated with hormone signaling in HR+ luminal cells, consistent with previous studies demonstrating that TNFSF11 (RANKL) and WNT control progesterone-mediated epithelial proliferation (*67*). Second, previous studies have shown that the fraction of apoptotic cells in the epithelium peaks between the late luteal and early follicular phases (*54*). Consistent with this, we identify subpopulations of HR+ and secretory luminal cells in the cycling premenopausal breast enriched for genes known to be upregulated during post-lactational involution (*51, 52*). Notably, we also observe upregulation of hypoxic gene signatures in multiple epithelial and stromal cell types that are highly correlated with hormone signaling in HR+ cells. A previous study identified these same pathways as highly enriched following involution in the mouse mammary gland. More importantly from the perspective of breast cancer risk, this “hypoxia/pro-angiogenic” signature identified breast cancers with increased metastatic activity (*68*), suggesting that these pathways support a permissive tumor microenvironment.

Finally, we find that paracrine signaling from HR+ cells to basal cells depends on both the per-cell transcriptional response of HR+ cells to hormones and the overall proportion of HR+ cells in the epithelium. Notably, prior pregnancy and obesity are specifically associated with a reduced risk of ER+/PR+ breast cancer in premenopausal women (*11, 12*), and our data support the idea that these biological variables lead to reduced paracrine signaling downstream of estrogen and progesterone via two distinct mechanisms. First, parity leads to a reduced per-cell hormone signaling response in HR+ luminal cells. Second, we identify a marked decrease in the ratio of HR+ cells relative to secretory luminal cells with increasing BMI. Both changes are associated with a reduced paracrine signaling response in basal/myoepithelial cells.

In summary, these results provide a comprehensive, systems-level view of the cellular and transcriptional variation within the disease-free human breast which profoundly affects the response to hormones and breast cancer risk. This single-cell analysis establishes a link between hormone signaling and tumor-promoting changes in cell state across multiple cell types. Furthermore, we identify tumor-protective changes in epithelial cell proportions and hormone responsiveness with pregnancy and increased BMI. As the breast is one of the only human organs that undergoes repeated cycles of morphogenesis and involution, this study serves as a roadmap to the cell state changes associated with hormone dynamics in the human breast. Finally, it provides a foundation for similar systems-level studies dissecting how the paracrine communication networks downstream of hormone signaling are altered during ER+/PR+ breast cancer progression.

## Acknowledgments

We thank Drs. Tom Norman and Jonathan Weissman for technical support and for generously providing access to equipment and computing resources. Sequencing was performed in the Center for Advanced Technology at UCSF. Tissue samples were provided by the Cooperative Human Tissue Network (CHTN), which is funded by the National Cancer Institute. Other investigators may have received specimens from the same subjects. Samples from the Susan G. Komen Tissue Bank at the IU Simon Cancer Center were used in this study. We thank contributors, including Indiana University who collected samples used in this study, as well as donors and their families, whose help and participation made this work possible. This research was supported in part by grants from the Department of Defense Breast Cancer Research Program (W81XWH-10-1-1023 and W81XWH-13-1-0221), NIH (U01CA199315 and DP2 HD080351-01), the NSF (MCB-1330864), and the UCSF Center for Cellular Construction (DBI-1548297), an NSF Science and Technology Center, to Z.J.G. Z.J.G is a Chan-Zuckerberg BioHub Investigator. L.M.M is a Damon Runyon Fellow supported by the Damon Runyon Cancer Research Foundation (DRG-2239-15).

## Author Contributions

L.M.M., R.J.W., and Z.J.G. conceived the project. L.M.M., J.A.C., R.J.W., C.S.M., and K.P. performed the sequencing experiments. C.S.M. generated aligned reads and barcode matrices. C.S.M and L.M.M. performed sample demultiplexing. P.G. coordinated sample acquisition and provided critical guidance for sample selection. L.M.M. performed fluorescent immunohistochemistry and RNA-FISH experiments. L.M.M. and J.A.C. performed flow cytometry experiments. L.M.M. performed histopathology on tissue sections. A.D.B. performed histopathological tissue analysis. L.M.M. analyzed and visualized the data. M.T. provided guidance in data analyses and computational approaches. T.T. and A.D.B. provided guidance in human breast biology. T.T., M.T., and Z.J.G. provided critical resources. T.A.D., M.T., T.T., and Z.J.G. supervised the project. L.M.M. and Z.G. wrote the manuscript. All authors reviewed and edited the manuscript.

## Materials and Methods

### Tissue samples and preparation

Reduction mammoplasty tissue samples were obtained from the Cooperative Human Tissue Network (CHTN, Vanderbilt University Medical Center, Nashville, TN) and Kaiser Permanente Northern California (KPNC, Oakland, CA). Core biopsy samples were provided by the Susan G. Komen Tissue Bank (KTB). Tissues were obtained as de-identified samples and all subjects provided written informed consent. When possible, medical reports or other patient data were obtained with personally identifiable information redacted. Use of breast tissue specimens to conduct the studies described above were approved by the UCSF Committee on Human Research under Institutional Review Board protocols 16-18865 and 10-01532. A portion of each sample was fixed in formalin and paraffin-embedded using standard procedures.

The remainder was dissociated mechanically and enzymatically to obtain epithelial-enriched tissue fragments. Tissue was minced, followed by enzymatic dissociation with 200 U/mL collagenase type III (Worthington CLS-3) and 100 U/mL hyaluronidase (Sigma H3506) in RPMI 1640 with HEPES (Corning 10-041-CV) plus 10% (v/v) dialyzed FBS, penicillin, streptomycin, amphotericin B (Lonza 17-836E), and gentamicin (Lonza 17-518) at 37 °C for 16 h. For KTB samples, the resulting cell suspension containing single cells and stroma was frozen and maintained at -180 °C until use. For reduction mammoplasty samples, the cell suspension was centrifuged at 400 x g for 10 min and resuspended in RPMI 1640 plus 10% FBS. Digested tissue fragments enriched for epithelial cells and closely-associated stroma were collected after serial filtration through 150 µm and 40 µm nylon mesh strainers. Following centrifugation, tissue fragments and filtrate were frozen and maintained at -180 °C until use.

### Dissociation to single cells

The day of sorting, epithelial-enriched tissue fragments from the 150 µm fraction, or total banked material for the KTB samples, were thawed and digested to single cells by trituration in 0.05% trypsin for 2 min, followed by trituration in 5 U/mL dispase (Stem Cell Technologies 07913) plus 1 mg/mL DNase I (Stem Cell Technologies 07900) for 2 min. Single-cell suspensions were resuspended in HBSS supplemented with 2% FBS, filtered through a 40 µm cell strainer, and pelleted at 400 x g for 5 min. The pellets were resuspended in 10 mL of complete mammary epithelial growth medium with 2% v/v FBS without GA-1000 (MEGM; Lonza CC-3150). Cells were incubated at 37 °C for 1 h, rotating on a hula mixer, to regenerate surface antigens.

### MULTI-seq sample barcoding

Single-cell suspensions were pelleted at 400 x g for 5 min and washed once with 10 mL mammary epithelial basal medium (MEBM; Lonza CC-3151). For each sample, one million cells were aliquoted, washed a second time with 200 μL MEBM, and resuspended in 90 μL of a 200 nM solution containing equimolar amounts of anchor lipid-modified oligonucleotides (LMOs) and sample barcode oligonucleotides in phosphate buffered saline (PBS). Following a 5-minute incubation on ice with anchor-LMO/barcode, 10 uL of 2 μM co-anchor LMO in PBS was added to each sample (for a final concentration of 200 nM), and wells were mixed by gentle pipetting and incubated for an additional 5 min on ice. Following incubation, cells were washed twice in 200 μL PBS with 1% BSA and pooled together into a single 15 mL conical tube containing 10 mL PBS/1% BSA. All subsequent steps were performed on ice.

### Sorting for scRNAseq

Cells were pelleted at 400 x g for 5 min and resuspended in PBS/1% BSA at a concentration of 1 million cells per 100 μL, and incubated with primary antibodies. Cells were stained with Alexa 488-conjugated anti-CD49f to isolate basal/myoepithelial cells, PE-conjugated anti-EpCAM to isolate luminal epithelial cells, and biotinylated antibodies for lineage markers CD2, CD3, CD16, CD64, CD31, and CD45 to remove hematopoietic (CD16/CD64-positive), endothelial (CD31-positive), and leukocytic (CD2/CD3/CD45-positive) lineages by negative selection (Lin-). Sequential incubation with primary antibodies was performed for 30 min on ice in PBS/1% BSA, and cells were washed with cold PBS/1% BSA. Biotinylated primary antibodies were detected with a streptavidin-Brilliant Violet 785 conjugate. After incubation, cells were washed once and resuspended in PBS/1% BSA plus 1 ug/mL DAPI for live/dead discrimination. Cell sorting was performed on a FACSAria II cell sorter. Live/singlet (DAPI-), luminal (DAPI-/Lin-/CD49f-/EpCAM+), basal/myoepithelial (DAPI-/Lin-/CD49f+/EpCAM-), or total epithelial (pooled luminal and basal/myoepithelial) cells were collected for each sample as specified in table S2 and resuspended in PBS/1% BSA at a concentration of 1000 cells/µL. For Batch 4, an aliquot of MULTI-seq barcoded cells were separately stained with biotinylated-CD45/streptavidin-Brilliant Violet 785 to enrich for immune cells, and sorted CD45+ cells were pooled with the Live/singlet fraction as specified in table S2.

Antibodies and dilutions used (µL/million cells) were as follows: FITC-EpCAM (1.5 µL; BD 550257, clone AD2), APC-CD49f (4 µL; Stem Cell Technologies 10109, clone VU1D9), Biotin-CD2 (8 µL; BioLegend 313636, clone GoH3), Biotin-CD3 (8 µL; BD 55325, clone RPA-2.10), Biotin-CD16 (8 µL; BD 55338, clone HIT3a), Biotin-CD64 (8 µL; BD 555526, clone 10.1), Biotin-CD31 (4 µL; Invitrogen MHCD31154, clone MBC78.2), Biotin-CD45 (1 µL; BioLegend 304004, clone HI30), BV785-Streptavidin (1 µL; BioLegend 405249).

### scRNAseq library preparation

cDNA libraries were prepared using the 10X Genomics Single Cell V2 (CG00052 Single Cell 3’ Reagent Kit v2: User Guide Rev B) or Single Cell V3 (CG000183 Single Cell 3’ Reagent Kit v3: User Guide Rev B) standard workflows as specified in table S2. Library concentrations were quantified using high sensitivity DNA Bioanalyzer chips (Agilent, 5067-4626) and Qubit dsDNA HS Assay Kit (Thermo Fisher Q32851). Individual libraries were sequenced on a lane of a HiSeq4500 or NovaSeq, as specified in table S2, for an average of ∼150,000 reads/cell.

### Expression library pre-processing

Cell Ranger (10x Genomics) was used to align sequences, filter data and count unique molecular identifiers (UMIs). Data were mapped to the human reference genome GRCh37 (hg19). The resulting sequencing statistics are summarized in table S2. For each experimental batch, the cellranger aggr pipeline (10X Genomics) was used to normalize read depth across droplet microfluidic lanes.

### Cell calling

For V2 experiments, cell-associated barcodes were defined using Cell Ranger. For V3/MULTI-seq experiments, cells were defined as barcodes associated with ≥600 total RNA UMIs and ≤20% of reads mapping to mitochondrial genes. We manually selected 600 RNA UMIs and 20% mitochondrial genes to exclude low-quality cell barcodes.

### MULTI-seq barcode library pre-processing

Raw barcode FASTQs were converted to barcode UMI count matrices as described previously (*16*). Briefly, FASTQs were parsed to discard reads where: 1) the first 16 bases of read 1 did not match a list of cell barcodes generated as described above, and 2) the first 8 bases of read 2 did not align with any reference barcode with less than 1 mismatch. Duplicated UMIs, defined as reads with the same cell barcode where bases 17-28 (V3 chemistry) of read 2 exactly matched, were removed to produce a final barcode UMI count matrix.

### Sample demultiplexing

Barcode UMI count matrices were used to classify cells using the MULTI-seq classification suite (*16*). In Batch 3, sample RM192 was poorly labeled for the lane of cells from the epithelial cell sort gate. Therefore, to reduce spurious doublet calls in this dataset, we manually set UMI counts which were <10 for this barcode to zero. For all experiments, raw barcode reads were log2-transformed and mean-centered, the top and bottom 0.1% of values for each barcode were excluded, and a probability density function (PDF) was constructed for each barcode. Next, all local maxima were computed for each PDF, and the negative and positive maxima were selected. To define a threshold between these two maxima, we iterated across 0.02-quantile increments and chose the quantile maximizing the number of singlet classifications, defined as cells surpassing the threshold for a single barcode. Multiplets were defined as cells surpassing two or more thresholds, and unlabeled cells were defined as cells surpassing zero thresholds. Unclassified cells were removed and the procedure was repeated until all remaining cells were classified.

To classify cells that were identified as unlabeled by MULTI-seq, we used the souporcell pipeline (*15*) to assign cells to different individuals based on single nucleotide polymorphisms (SNPs). For each dataset, we set the number of clusters (k) to the total number of samples in that experiment. To avoid local minima, souporcell restarts clustering multiple times and takes the solution that minimizes the loss function. For Batch 3, we chose the number of restarts that produced less than a 1.5% misclassification rate between MULTI-seq and souporcell singlet sample classifications (Live singlet: 30 restarts/1.2% mismatch rate; Epithelial: 75 restarts/1.5% mismatch rate). Souporcell classification performed more poorly across parameters for Batch 4 (Live singlet plus CD45+: 50 restarts/8.1% mismatch rate, 75 restarts/4.8% mismatch rate; Epithelial: 50 restarts/8.6% mismatch rate, 75 restarts/14.9% mismatch rate, 100 restarts/4.1% mismatch rate). Therefore, for these datasets we used sample classifications that were consistent across two restarts (Pooled live singlet/ CD45+: consistent calls across 50 and 75 restarts/0.4% overall mismatch rate; Epithelial: consistent calls across 50 and 100 restarts/1% overall mismatch rate) to identify high-confidence singlets.

### Quality control, dataset integration, and cell type identification using Seurat

Cell type identification was performed using the Seurat package (version 3.1.5) in R (*69*). To identify and remove doublets from the same sample that would not be identified by MULTI-seq or souporcell, we filtered each lane to remove cells with greater than 20% of reads mapping to mitochondrial genes and ran DoubletFinder (version 2.0) on each data subset (*70*), using parameters identified by the ‘paramSweep_v3’ function. Aggregated data for singlet cells for each batch was filtered to remove cells that had fewer than 200 genes and genes that appeared in fewer than 3 cells. Cells with a Z score of 4 or greater for the total number of genes expressed were presumed to be doublets and removed from analysis. The remaining cells were log transformed and scaled to a total of 1e4 molecules per cell, and the top 2000 most variable genes based on variance stabilizing transformation were identified for each batch (*71*). Data from all four batches were integrated using the standard workflow and default parameters from Seurat v3 (*69*). This data integration workflow identifies pairwise correspondences between cells across datasets and uses these anchors to transform datasets into a shared expression space. Following dataset integration, the resulting batch-corrected expression matrix was scaled, and principal component (PC) analysis was performed using the identified integration genes. The top 28 statistically significant PCs as determined by visual inspection of elbow plots were used as an input for UMAP visualization and k-nearest neighbor (KNN) modularity optimization-based clustering using Seurat’s FindNeighbors and FindClusters functions.

### Quantification of sample-to-sample heterogeneity: cluster entropy and similarity score

To measure how well-mixed cells from different samples were across cell type clusters, we quantified the normalized relative cluster entropy for our dataset, weighted by cluster size (*72*). A cluster entropy value of 1 represents complete intermixing of samples across clusters. To measure transcriptional variation in cell state within cell types between cells from the same versus different batches and/or samples, we measured the pairwise alignment between each sample/batch (*73*), where batches consisted of sets of samples processed on the same day (table S2). This “similarity score” examines the local neighborhood of each cell in a particular sample/batch, asks how many of its k nearest neighbors belong to a second sample/batch, and averages this over all cells. We chose k to be 1% of the total number of cells within a cluster. The result was normalized by the expected number of cells from each sample/batch. Notably, for repeat measurements, samples run across multiple batches were highly similar.

### Testing for changes in cell type proportions and predictive modeling

We modeled the detected number of each cell type in each sample as a random count variable using a quasi-Poisson process to allow for overdispersion, with the condition being tested (e.g. parity, BMI, obesity) as a predictor and the total number of detected epithelial or luminal cells in each sample as an offset variable (*74*). To account for uncertainty due to variable numbers of profiled cells in each sample, we used bootstrap resampling to estimate confidence intervals associated with detection of each cell type (*75*). Results from 1000 bootstrap replicates were pooled using the mice::pool function in R, and the model was fit using a quasi-Poisson generalized linear model from the ‘stats’ R package. Tests for statistical significance were performed using a Wald test on the regression coefficient. Multiple hypothesis correction was controlled using the false discovery rate. For the Komen Tissue Bank (KTB) data set, a quasi-Poisson model was trained on the reduction mammoplasty cohort as described above, and the ‘predict’ function in the ‘stats’ R package was used to predict the proportion of HR+ luminal cells in the KTB samples based on BMI.

### PC analysis within HR+ luminal cells

To perform principal component analysis on HR+ luminal cells, we subset out this cluster from the integrated dataset and repeated the standard workflow from Seurat v3 to identify integration genes specific to this cell type. The resulting batch-corrected expression matrices were scaled, and PC analysis was performed using the identified integration genes.

### Non-negative matrix factorization of individual cell types

To identify gene expression signatures, or “metagenes” within individual cell types, we subset out raw counts data from each of the four most abundant clusters (HR+ luminal cells, secretory luminal cells, basal/myoepithelial cells, and fibroblasts) and performed matrix factorization. We chose to perform matrix factorization independently on each cell type rather than on the combined dataset, as preliminary analyses demonstrated that the number of metagenes identified for each cell type was highly dependent on the relative sizes of each cluster in the combined dataset. To account for batch differences, we used the LIGER package in R to perform integrative NMF (*37, 38*), and performed all subsequent analyses on shared, rather than batch-specific, metagenes. To avoid identification of gene signatures dominated by highly-expressed transcripts, we normalized the raw counts matrix for each cell based on its total expression, multiplied by a scale factor of 1e4, and log-transformed and scaled the result without centering. The resulting dataset was decomposed using the standard workflow and default parameters from LIGER. To estimate the optimum choice of rank *K* (i.e. number of NMF components) for each cell type, we used the suggestK function in the LIGER package to calculate the Kullback-Leibler (KL) divergence of metagene loadings across a range of *K* values, and identified the elbow point on this curve.

### Jensen-Shannon distance to quantify sample-to-sample variability in hormone signaling

To quantify variation in expression of the “hormone signaling” metagene in HR+ luminal cells (HR+ metagene 8), we performed the following workflow. First, we used the cell loadings across HR+ metagene 8 for each sample to compute kernel density estimations using the ‘density’ function in the ‘stats’ R package. We excluded sample RM172 from this analysis as it had fewer than 50 HR+ luminal cells; thus, the resulting kernel density estimation was highly sensitive to individual outliers. Second, we used the ‘JSD’ function in the ‘philentropy’ R package (*76*) to measure the pairwise Jensen-Shannon divergence between samples. Third, we converted this to a distance metric (Jensen-Shannon Distance, JSD) by taking the square root and performed hierarchical clustering using the ‘hclust’ function in the ‘stats’ R package, using ‘ward.D2’ linkage. The similarity between samples was plotted on a heatmap as (1-JSD).

### Metagene network analysis

To identify sets of gene expression programs that co-varied across samples, we first decomposed each cell type into a set of distinct gene expression programs, or “metagenes”, using NMF as described above. We then quantified the average expression of each metagene in each sample and constructed a weighted network of coordinated gene expression programs based on the pair-wise Pearson correlations between metagenes. To account for correlations driven by outlier samples, we used bootstrap resampling to estimate confidence intervals associated with each correlation coefficient. The resulting Pearson correlation matrix was transformed into a weighted adjacency matrix by setting all Pearson correlation coefficients less than 0.5 or with p-values greater than 0.05 to zero. Finally, we identified modules of highly correlated gene expression programs using the infomap community detection algorithm in the ‘igraph’ package in R (*45*). We chose this flow-based community detection algorithm in order to maximize information flow within clusters. Results using the modularity-based Louvain clustering algorithm were identical except that a small community consisting of three metagenes was merged with the “involution” module.

### Gene set enrichment analysis

To identify marker genes statistically associated with each metagene, we used multiple least squares regression of normalized (z-scored) gene expression against the cell loading matrix for each metagene (*77*). This results in a vector of regression coefficients representing the strength of the relationship between expression of a particular metagene and scaled expression of each gene. The resulting ranked gene lists were analyzed by gene set enrichment analysis, using the ‘fgsea’ package in R (*78*).

### Fluorescent Immunohistochemistry

For immunofluorescent staining, formalin-fixed paraffin-embedded tissue sections were deparaffinized and rehydrated using standard methods. Endogenous peroxides were blocked using 3% hydrogen peroxide in PBS, and antigen retrieval was performed in 0.1 M citrate buffer pH 6.0. Sections were blocked for 5 min at room temperature using Lab Vision Ultra-V block (Thermo TA-125-UB) and rinsed with TNT wash buffer (1X Tris-buffered saline with 5 mM Tris-HCl and 0.5% TWEEN-20). Primary antibody incubations were performed for 1 hour at room temperature or overnight at 4°C. Sections were washed three times for 5 min each with TNT wash buffer, incubated with Lab Vision UltraVision LP Detection System HRP Polymer (Thermo Fisher TL-060-HL) for 15 min at room temperature, washed, and incubated with one of three colors of tyramide signal amplification amplification (TSA) reagent at a 1:50 dilution. After TSA, antibody complexes were removed by boiling in citrate buffer, followed by blocking and incubation with additional primary antibodies as above. Finally, sections were rinsed with deionized water and mounted using Vectashield HardSet Mounting Media with DAPI (Vector H-1400). Immunofluorescence was analyzed by spinning disk confocal microscopy using a Zeiss Cell Observer Z1 equipped with a Yokagawa spinning disk and running Zeiss Zen Software.

Antibodies, TSA reagents, and dilutions used are as follows: p63 (1:2000; CST 13109, clone D2K8X), KRT7 (1:4000; Abcam AB68459, clone EPR1619Y), KRT23 (1:2000; Abcam AB156569, clone EPR10943), ER (1:4000; Thermo Scientific RMM-9101-S, clone SP1), PR (1:3000; CST 8757, clone D8Q2J), TCF7 (1:2000; CST 2203, clone C63D9), FITC-TSA (2 min; Perkin Elmer NEL701A001KT), Cy3-TSA (3 min; Perkin Elmer NEL744001KT), Cy5-TSA (7 min; Perkin Elmer NEL745E001KT).

### Morphometric analysis and geometric modeling

Formalin-fixed paraffin-embedded tissue sections were immunostained for the pan-luminal marker KRT7, counterstained with DAPI and imaged as described above. Images containing lobular tissue were acquired randomly, and the area and perimeter of the KRT7-positive luminal layer of each acinus was analyzed in ImageJ. To reduce noise and remove small gaps in KRT7 fluorescence, we applied a closing filter from the MorphoLibJ plugin with a 2-pixel (1.33 μm) radius disk (*79*). The resulting image was smoothed by applying a Gaussian filter with sigma 5 pixels (3.33 μm), and binarized using the default thresholding algorithm in ImageJ. Finally, individual acini with visible lumens were manually selected and the area (*A*), perimeter (*P*), and circularity of the KRT7-positive region was measured for each structure. To estimate the average diameter (*d*) and luminal thickness (*w*) of each acinus, we used area and perimeter measurements to fit a circle containing a hollow lumen to each structure. Based on these results, we implemented a geometric model in which each acinus was represented as a hollow circle with shell thickness that was linearly related to diameter (*d*). Since basal cells form a monolayer along the luminal surface, we represented the space available for basal cells as the outer perimeter of the luminal layer, and the space available for luminal cells as the area of the luminal layer. To estimate the linear relationship between *w* and *d*, we performed linear regression analysis using measurements from all structures with a circularity greater than 0.75 (n = 55 acini from 15 samples).

### Pseudo-bulk differential gene expression analysis

To identify genes differentially expressed between samples from parous and nulliparous or obese and non-obese individuals in specific cell types, we constructed pseudo-bulk datasets consisting of the summed raw read counts across all single HR+ luminal cells or basal/myoepithelial cells for each batch and sample. We restricted our analysis to samples/batches that had at least 100 cells of the cell type of interest. Each dataset was then randomly down-sampled to the lowest library size, and differential expression analysis was performed using DESeq2 (version 1.18.1) to test for genes differentially expressed between samples from obese (BMI ≥ 30) and non-obese (BMI < 30) or parous and nulliparous individuals, using batch as a covariate (*80*). As certain samples were sequenced across more than one batch (table S2), replicates of the same sample from different batches were combined using the collapseReplicates function. False discovery rate corrected p-values were calculated using the Benjamini-Hochberg procedure (*81*).

### RNA FISH analysis of ESR1 transcripts

Combined RNA FISH and immunofluorescence analysis of estrogen receptor transcript (RNAscope Probe Hs-ESR1; ACD 310301) and protein (anti-ER; Thermo RMM-9101-S, clone SP1) was performed using the RNAscope in situ hybridization kit (RNAscope Multiplex Fluorescent Reagent Kit V2, ACD 323100) according to the manufacturer’s instructions and fluorescent immunohistochemistry protocol outlined above with the following modifications. Immunostaining for ER was performed prior to *in situ* hybridization, using the hydrogen peroxide and antigen retrieval solutions supplied with the RNAscope kit and the mildest recommended conditions. After ER immunostaining and tyramide signal amplification, *in situ* hybridization for ESR1 was performed according to the manufacturer’s instructions, followed by immunostaining for KRT7 as described above. For all RNA FISH experiments, we used positive (PPIB) and negative controls (DAPB) to verify staining conditions and probe specificity.

### Data availability

Submission of raw gene expression and barcode count matrices to the Gene Expression Omnibus is in process. For inquiries about data, code, or materials, contact authors.

## Supplemental Figures and Tables

Tables S1-S7

Figures S1-S16

## Supplemental Figures and Tables

**Table S1**. Donor information for reduction mammoplasty samples and list of samples used for scRNAseq, FACS, and immunostaining experiments.

**Table S2**. Summary statistics for single-cell RNA sequencing of twenty-eight reduction mammoplasty samples (RM) and seven Komen Tissue Bank samples (KTB).

**Table S3**. Multiple linear regression analysis of the percentage of basal cells in the epithelium as measured by flow cytometry analysis.

**Table S4**. Association of the 20 highest-loading genes in PC1 for HR+ luminal cells with estrogen signaling, progesterone signaling, or the luteal phase of the menstrual cycle.

**Table S5**. Canonical hormone-responsive genes differentially expressed in HR+ luminal cells between parous and nulliparous samples.

**Table S6**. Multiple linear regression analysis of the basal paracrine response (metagene 10) in response to three predictors: HR+ cell hormone signaling (HR+ metagene 8), the frequency of HR+ cells in the epithelium, and an interaction term representing the combined effects of HR+ signaling and frequency (Signaling × Frequency)

**Table S7**. Genes differentially expressed in basal/myoepithelial cells between parous versus nulliparous samples or obese (BMI ≥ 30) versus non-obese (BMI < 30) samples.

**Fig. S1.**
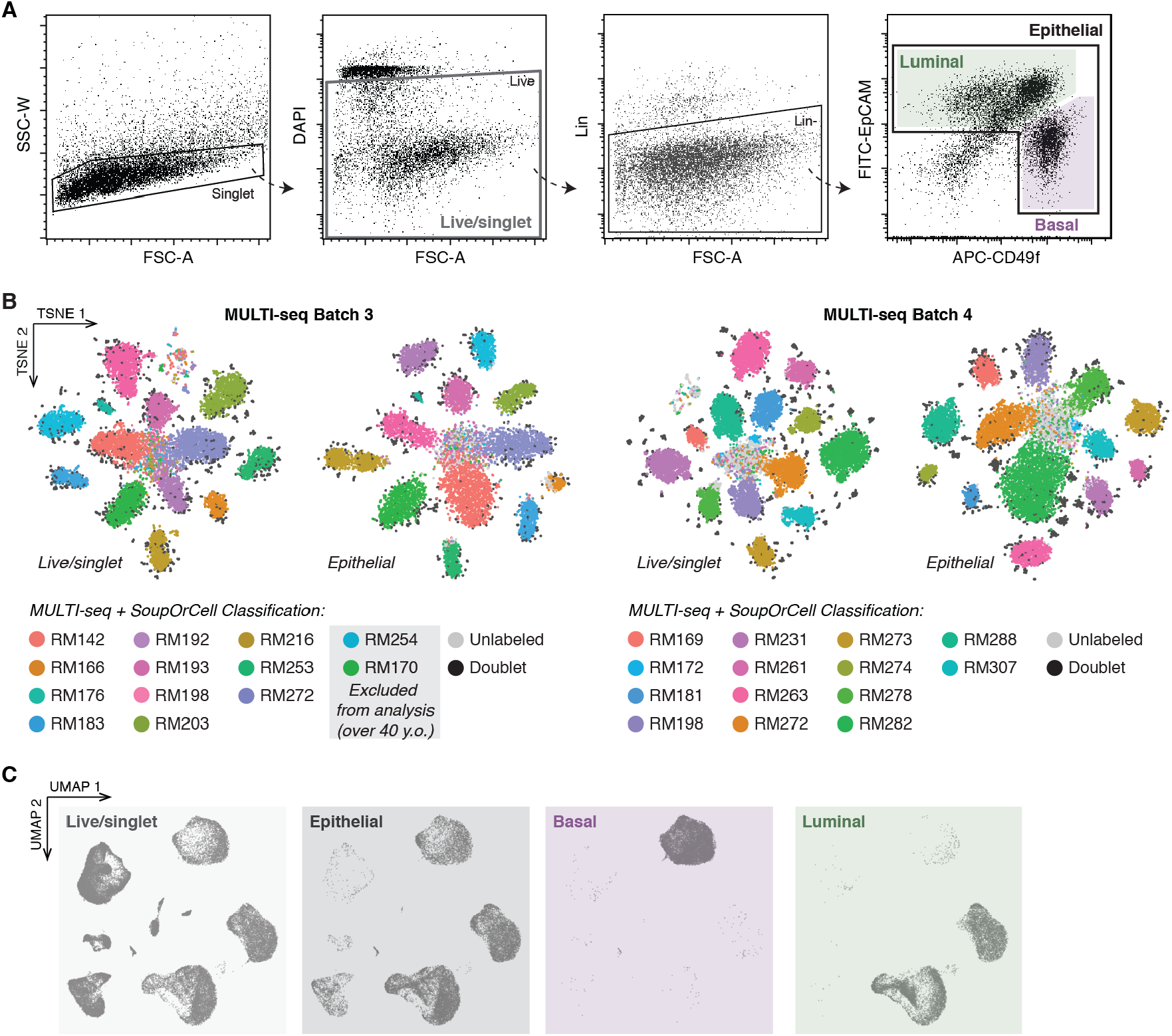
Sorting strategy and MULTI-seq barcoding for scRNAseq experiments. (A) FACS plots depicting sort gates used for sequencing. (B) TSNE dimensionality reduction of the normalized barcode count matrices and final sample classification for MULTI-seq experiments (Batches 3 and 4). (C) UMAP dimensionality reduction of the combined data from twenty-eight samples for each sort population.

**Fig. S2.**
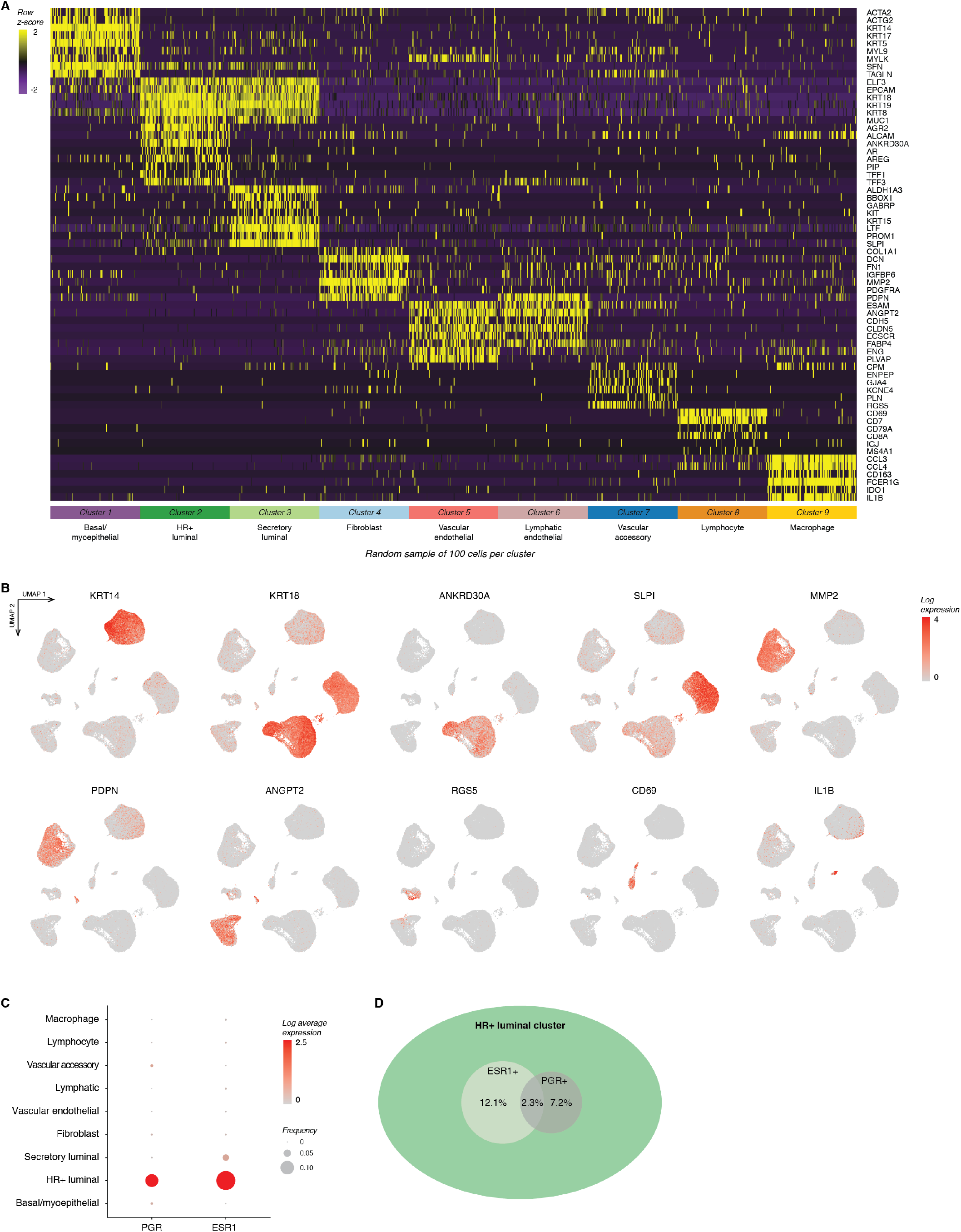
Marker analysis of cell type clusters for scRNAseq experiments. (A) Heatmap highlighting marker genes used to identify each cell type. For visualization purposes, we randomly selected 100 cells from each cluster. (B) UMAPs depicting expression of selected markers in log normalized counts. (C) Dot plot depicting the log normalized average and frequency of ESR1 and PGR expression across cell type clusters. (D) Venn diagram highlighting the frequency of ESR1 and PGR expression and percent overlap in the HR+ luminal cell cluster.

**Fig. S3.**
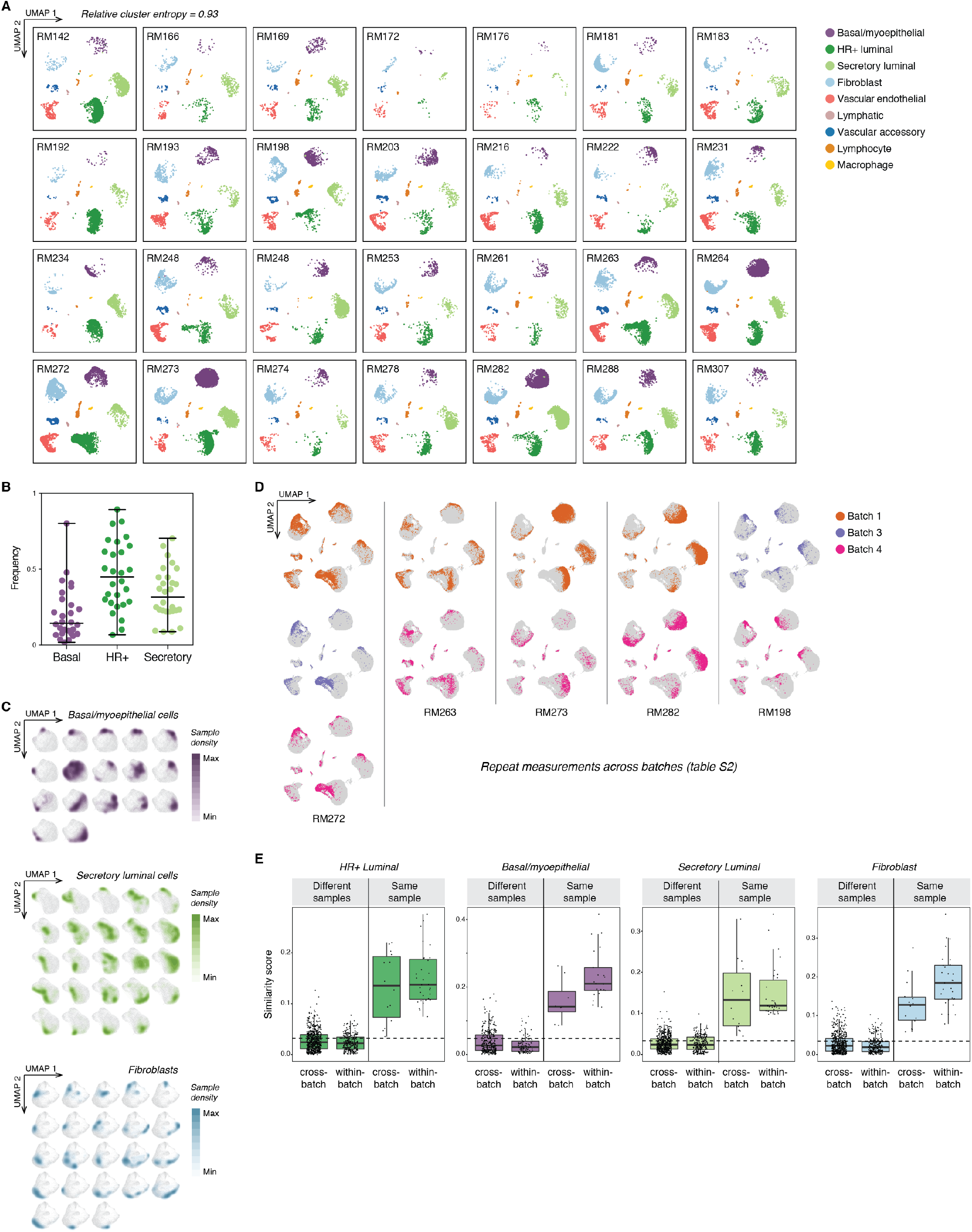
Quantification of inter-sample variability in the breast. (A) UMAP for each sample highlighting cell types identified by unsupervised clustering. Cells from different individuals are represented across all clusters (cluster entropy = 0.93, *methods*). (B) Quantification of the proportion of epithelial cells (basal/myoepithelial; HR+ luminal; secretory luminal) in each sample, with the cross-sample median and range for each cell type (n = 28 samples). (C) Density plots highlighting the transcriptional cell state of basal/myoepithelial cells, secretory luminal cells, or fibroblasts from samples with at least 100 cells in each cluster. (D) UMAP of samples that were run as repeat measurements across multiple batches, highlighting cells from each batch. See table S2 for sample and batch information. (E) Quantification of the pairwise alignment—or “similarity score”—between cells from the same or different sample and batch for the indicated cell types. See table S2 for sample and batch information. The dashed line represents the expected similarity score for random mixing.

**Fig. S4.**
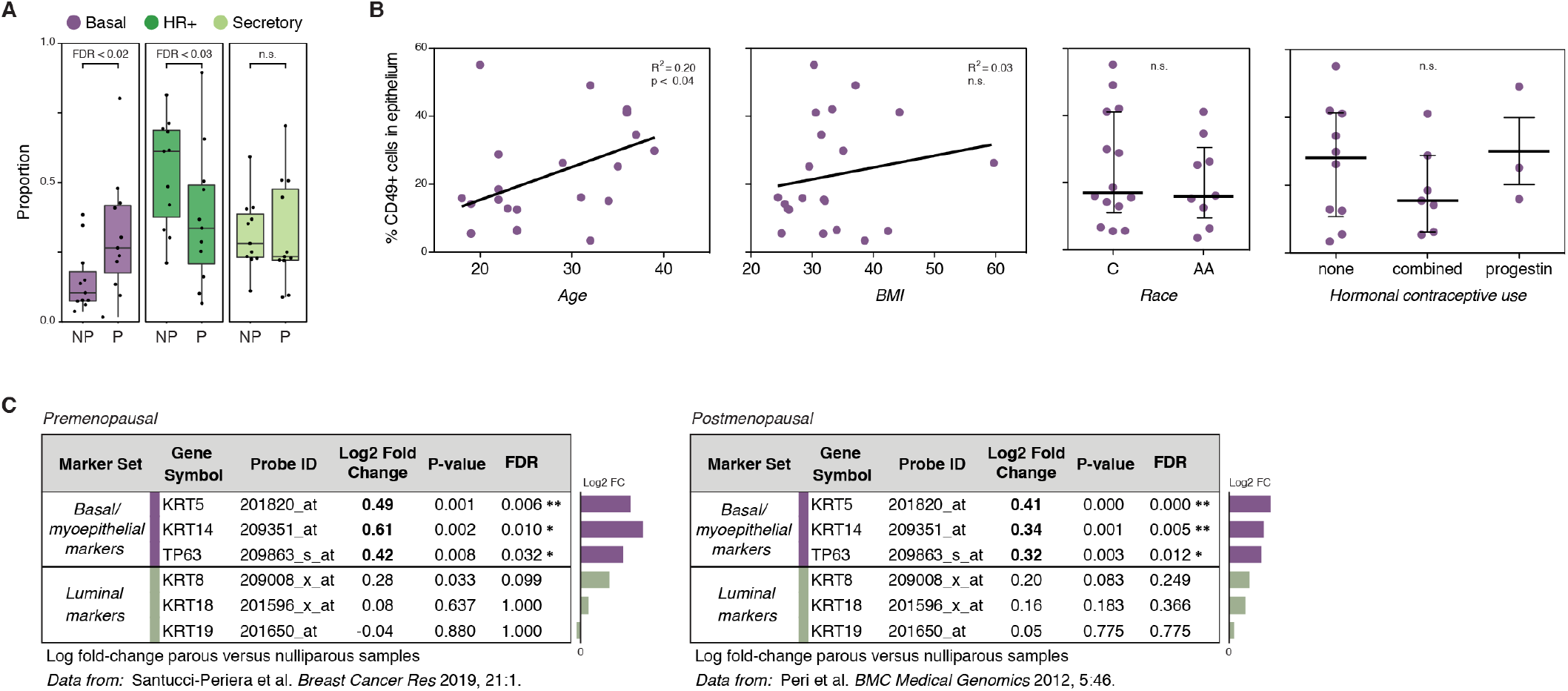
Prior pregnancy is associated with changes in epithelial cell proportions. (A) Quantification of the proportion of basal/myoepithelial cells (Basal), HR+ luminal cells (HR+), and secretory luminal cells (Secretory) in the mammary epithelium of nulliparous (NP) versus parous (P) samples, as identified by scRNAseq clustering (n = 28 samples; Wald test). (B) Quantification of the percentage of EpCAM^−^/CD49f^+^ basal cells identified by FACS analysis versus age (n = 23; R^2^ = 0.20; p < 0.04, Wald test), body mass index (n = 21; R^2^ = 0.03; p = 0.44, Wald test), race (Caucasian = C, African American = AA; n = 23; p = 0.55, Mann-Whitney test), or hormonal contraceptive use (n = 23; p = 0.50, Kruskal-Wallis test). (C) Microarray differential expression analysis for selected genes from Santucci-Periera *et al*. and Peri *et al*. (*20, 21*).

**Fig. S5.**
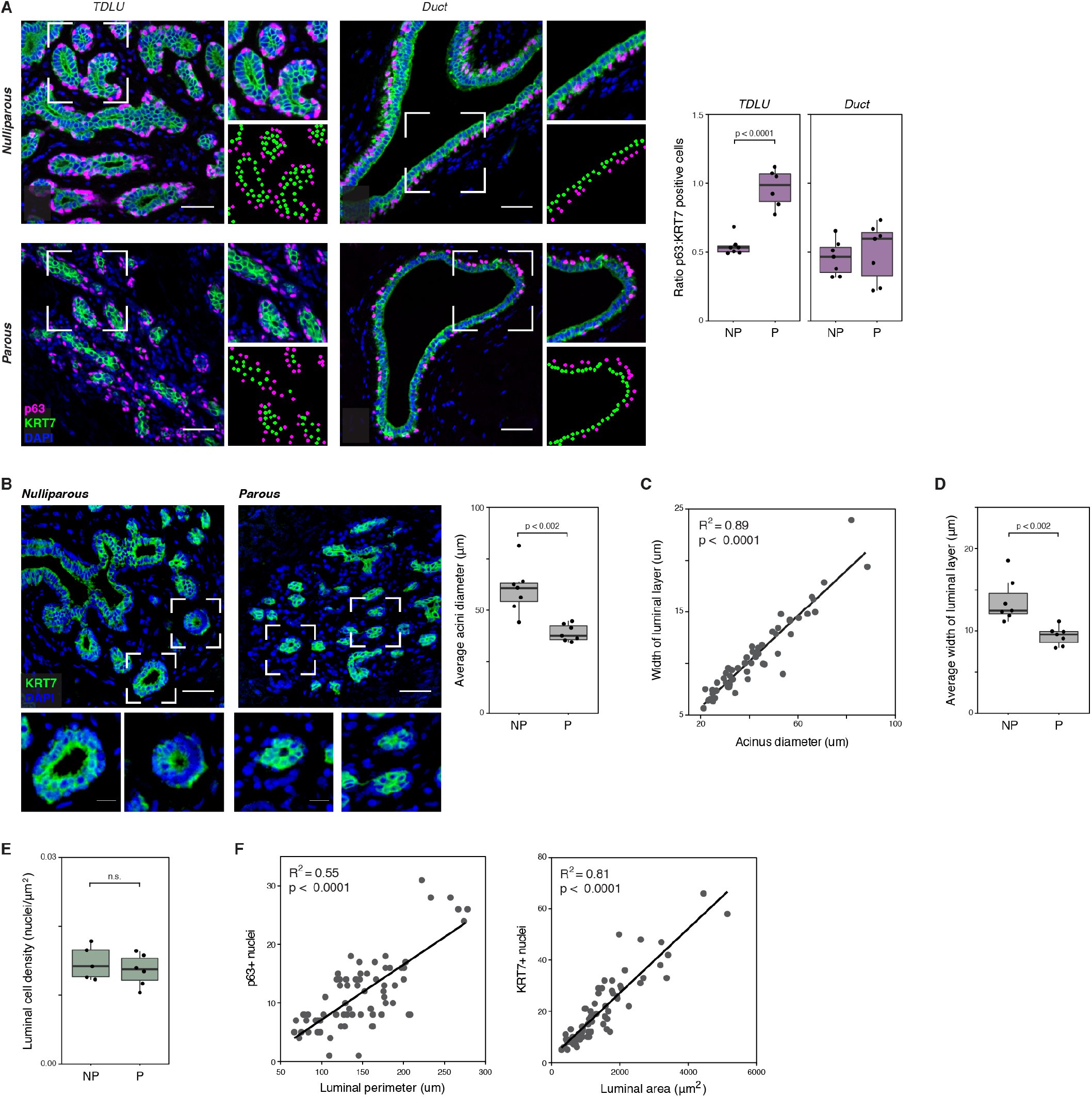
Prior pregnancy is associated with changes in epithelial architecture. (A) Immunostaining for the basal/myoepithelial marker p63 and pan-luminal marker KRT7, and quantification of the ratio of p63+ myoepithelial cells to KRT7+ luminal cells in the terminal ductal lobular units (TDLUs) or ducts for parous (P) versus nulliparous (NP) samples (n = 14 samples; Mann-Whitney test). Scale bars 50 µm. Inset scale bars 15 µm (B) Quantification of the average acinar diameter in TDLUs from nulliparous (NP) versus parous (P) samples (n = 14 samples; p < 0.002, Mann-Whitney test). (C) Linear regression analysis of the width of the luminal layer versus acinus diameter for individual acini with circularity greater than 0.75 (n = 56 acini from 15 samples; R^2^ = 0.89, p < 0.0001, Wald test). (D) Quantification of the average thickness of the luminal layer in acini from TDLUs in nulliparous (N) versus parous (P) samples (n = 14 samples; p < 0.002, Mann-Whitney test). (E) Quantification of the average luminal cell density (nuclei per μm^2^ of luminal area) in acini from TDLUs in nulliparous (NP) versus parous (P) samples (p = 0.43, Mann-Whitney test). (F) *Left:* Linear regression analysis of the perimeter of the luminal layer versus the number of p63+ basal cells for individual acini (n = 72 acini from 13 samples; R^2^ = 0.55, p < 0.0001, Wald test). *Right:* Linear regression analysis of the area of the luminal layer versus the number of KRT7+ luminal cells for individual acini (n = 72 acini from 13 samples; R^2^ = 0.81, p < 0.0001, Wald test).

**Fig. S6.**
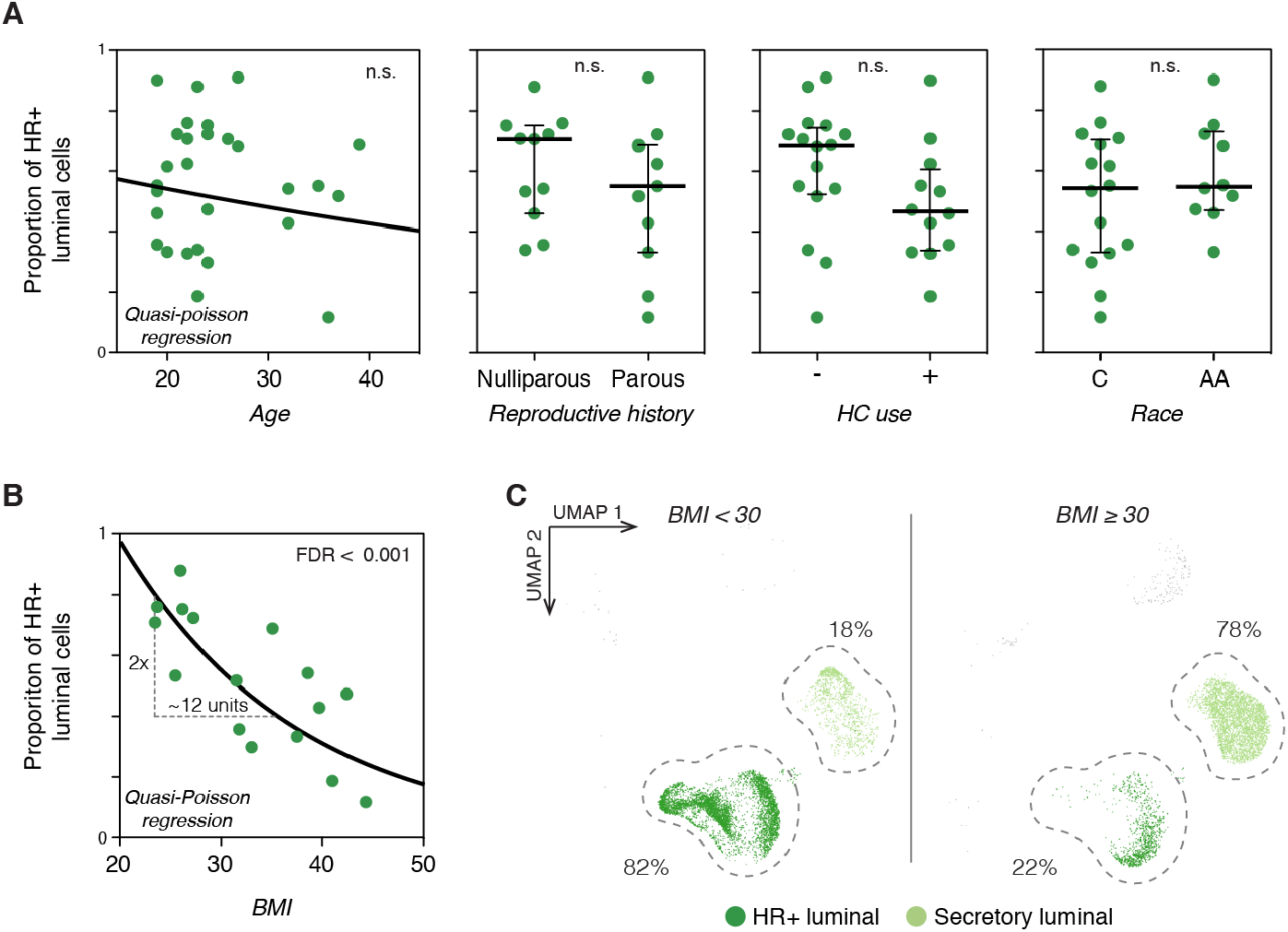
The proportion of HR+ luminal cells is reduced in obese women and does not vary with other discriminating factors. (A) Proportion of HR+ luminal cells in each sample (dots) stratified by age, reproductive history, hormonal contraceptive (HC) use, or race (C = Caucasian, AA = African American; Wald test). (B) Quasi-Poisson regression model of the proportion of HR+ cells in the luminal compartment as a function of BMI (FDR < 0.001, Wald test). (C) UMAP plot of sorted luminal cells from non-obese (BMI < 30) and obese (BMI ≥ 30) samples, highlighting hormone-responsive (HR+) and secretory luminal cells.

**Fig. S7.**
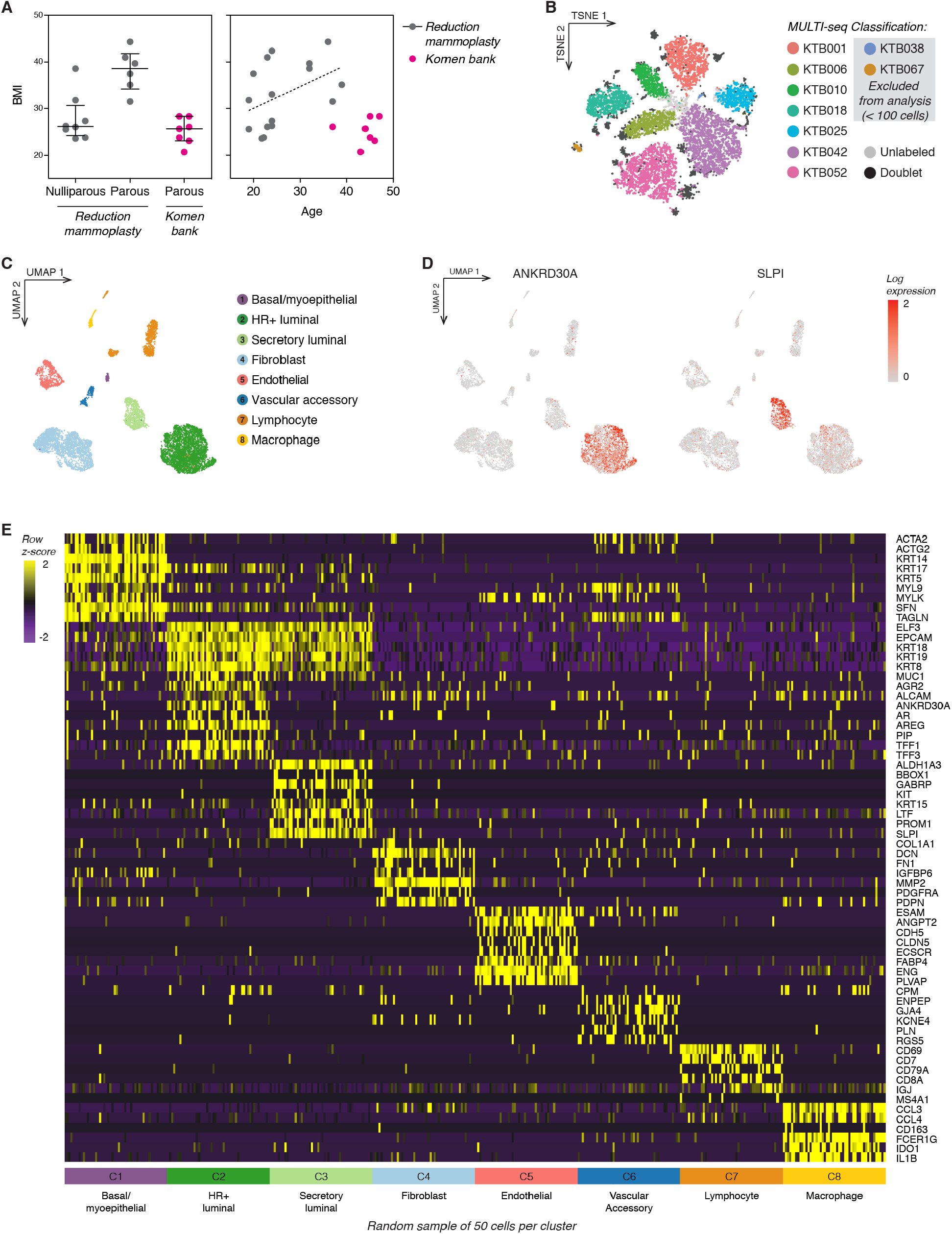
Summary of scRNAseq analysis of samples from the Komen Tissue Bank. (A) Scatter plots highlighting differences in body mass index (BMI), reproductive history, and age between the Komen Tissue Bank (KTB) and reduction mammoplasty cohorts (see also table S1). Trendline depicts the positive association of BMI with age in the reduction mammoplasty cohort. (B) TSNE dimensionality reduction of the normalized barcode count matrices and final sample classification for MULTI-seq barcoding. (C) UMAP dimensionality reduction and unsupervised clustering of the combined data from seven KTB samples identifies the major epithelial and stromal cell types in the breast. (D) UMAPs depicting expression of selected markers in log counts. (E) Heatmap highlighting marker genes used to identify each cell type. For visualization purposes, we randomly selected 50 cells from each cluster.

**Fig. S8.**
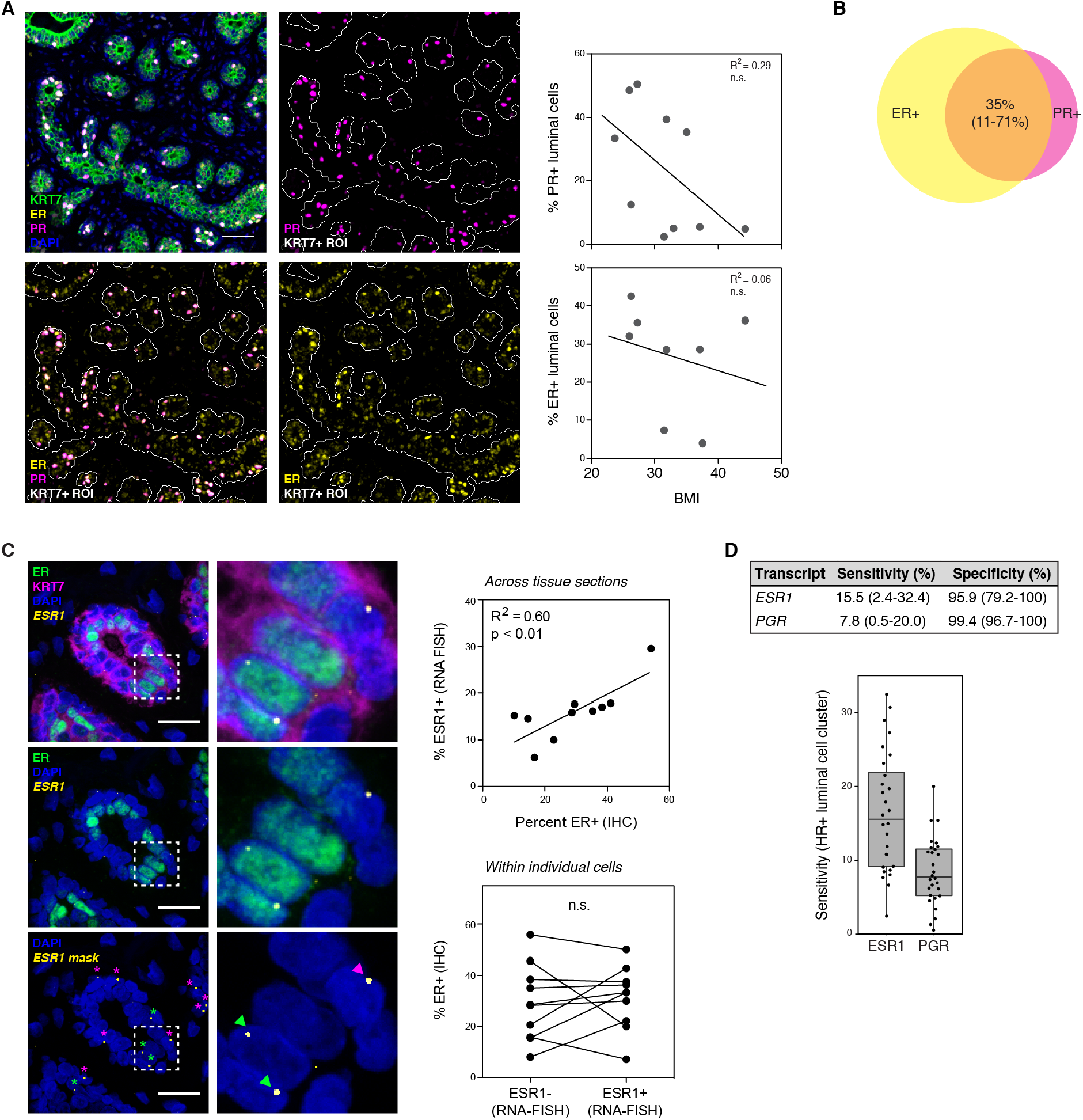
Hormone receptor expression is highly variable. (A) *Top:* Co-immunostaining of PR and KRT7 and linear regression analysis of the percentage of PR+ luminal cells versus BMI (n = 10 samples; R^2^ =0.29, p = 0.11, Wald test). *Bottom:* Co-immunostaining of ER and KRT7 and linear regression analysis of the percentage of ER+ luminal cells versus BMI (n = 8 samples; R^2^ =0.06, p = 0.56, Wald test). Scale bars 50 µm. (B) Venn diagram highlighting the average percent overlap between ER and PR as measured by immunostaining (n = 5 samples, range = 11-71%). (C) Multiplexed *in situ* hybridization of estrogen receptor transcript (ESR1) and immunostaining for estrogen receptor protein (ER) and KRT7. Scale bars 25 µm. *Right:* Plots depicting the expression of ESR1 and ER across multiple tissue sections (R^2^ = 0.6, p < 0.01, Wald test) or within individual cells (p = 0.63, Wilcoxon matched pairs signed-rank test). (D) Table and bar plot depicting the sensitivity and specificity for ESR1 or PGR transcript expression in the HR+ luminal cell versus secretory luminal cell cluster based on scRNAseq analysis.

**Fig. S9.**
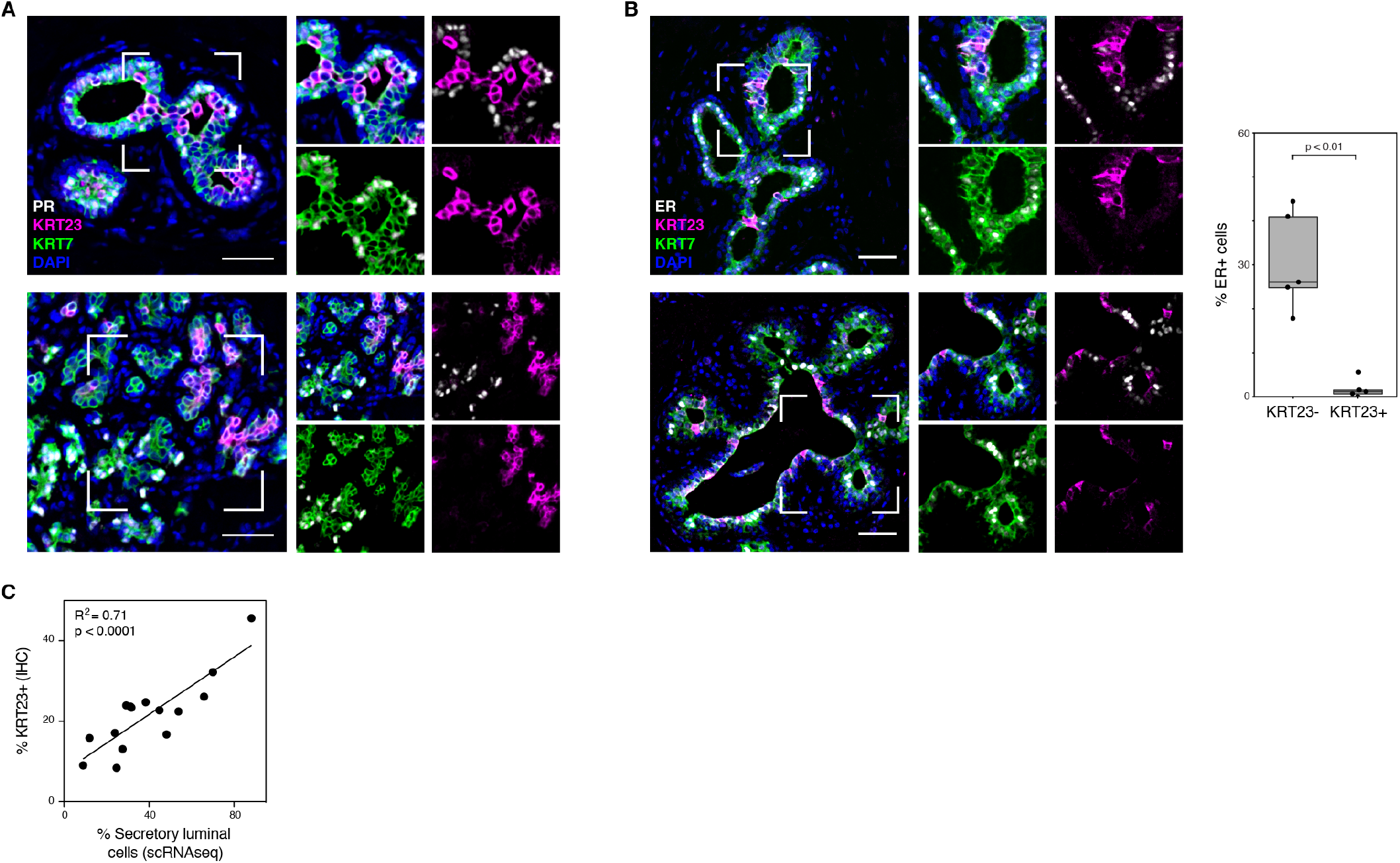
Keratin 23 is a specific marker of cells in the secretory luminal cell lineage. (A) Representative images of co-immunostaining of PR, KRT23, and the pan-luminal marker KRT7. (B) Co-immunostaining of ER, KRT23, and the pan-luminal marker KRT7 and quantification of the percentage of ER+ cells within the KRT7+/KRT23- and KRT7+/KRT23+ luminal cell populations (n = 5 samples; p < 0.01 Mann-Whitney test). Scale bars = 50 µm. (C) Linear regression analysis of the percentage of luminal cells in the secretory lineage identified by scRNAseq clustering versus the percentage of KRT23+ luminal cells identified by immunostaining (n = 15 samples; R^2^ =0.71, p < 0.0001, Wald test).

**Fig. S10.**
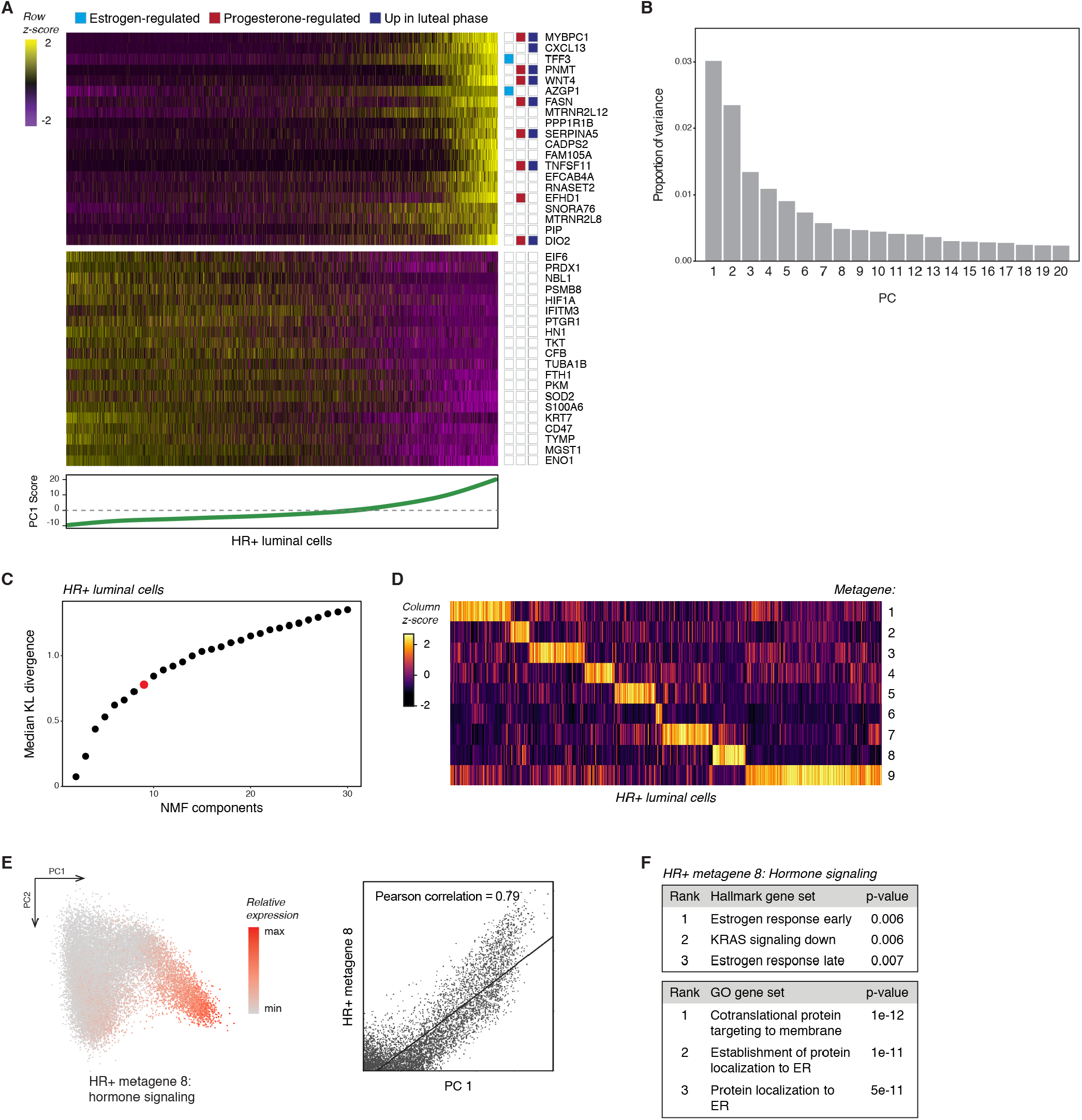
Matrix decomposition analysis of HR+ luminal cells. (A) Heatmap highlighting the 20 genes with the highest (top) and lowest (bottom) loadings in PC1, annotated by their association with estrogen signaling, progesterone signaling, or the luteal phase of the menstrual cycle. HR+ luminal cells are ordered by their cell loadings in PC1. (B) Barchart depicting the proportion of variance explained by each of the top 20 principal components. (C) Parameter selection for non-negative matrix factorization based on KL divergence (*methods*). (D) Heatmap of cell loadings across each metagene for HR+ luminal cells. (E) PCA plot of HR+ luminal cells depicting the relative expression of HR+ metagene 8. (F) Gene set enrichment analysis of HR+ cell metagene 8, showing the top pathways identified from the Molecular Signatures Database Hallmark and GO gene sets.

**Fig. S11.**
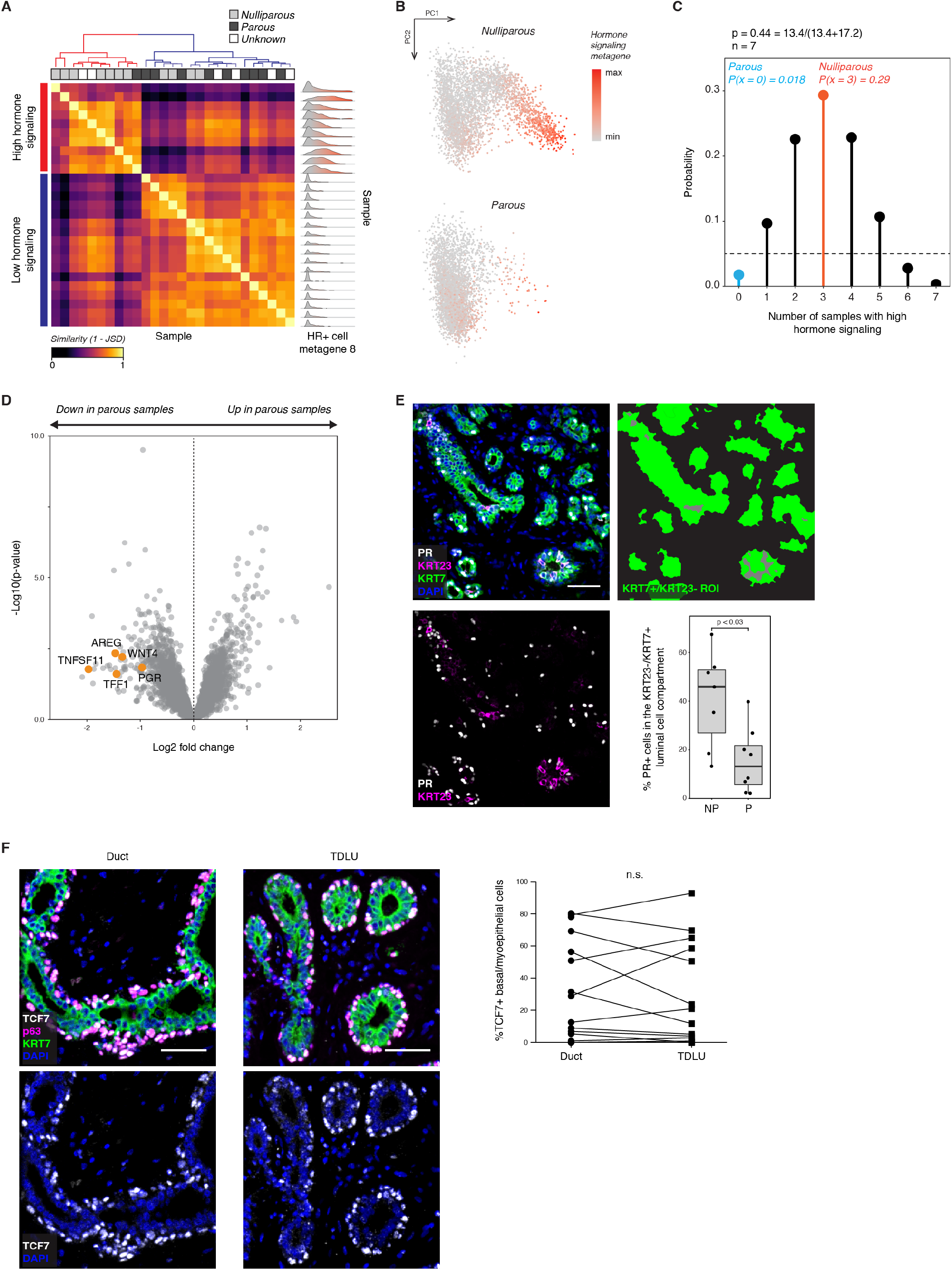
Parity is associated with a decrease in the per-cell hormone signaling response of HR+ luminal cells. (A) Heatmap showing the similarity between each sample’s single-cell expression distribution across HR+ cell metagene 8, measured as (1 - Jensen-Shannon distance). Hierarchical clustering identifies two sets of samples representing high or low expression of the “hormone signaling” metagene (ward D2). (B) PCA plot of HR+ luminal cells in nulliparous or parous women depicting expression of HR+ cell metagene 8. (C) Binomial probability distribution for the number of samples with high hormone signaling. The binomial probability of high hormone signaling is modeled as the average length of the luteal phase of the menstrual cycle, in days, divided by the average total length of the menstrual cycle (P = 0.44) (*40*). (D) Volcano plot highlighting the differential expression of canonical hormone-responsive genes between parous and nulliparous samples in HR+ luminal cells. (E) Immunostaining for PR, KRT23, and KRT7, and quantification of the percentage of PR+ cells within the KRT23-/KRT7+ luminal cell compartment for nulliparous (NP) versus parous (P) samples (n=15 samples; p < 0.03, Mann-Whitney test). (F) Immunostaining for p63, TCF7, and KRT7 in ducts versus TDLUs, and quantification of the percentage of TCF7+ cells within the p63+ basal/myoepithelial cell compartment (n = 14 samples; p = 0.64, Wilcoxon signed-rank test).

**Fig. S12.**
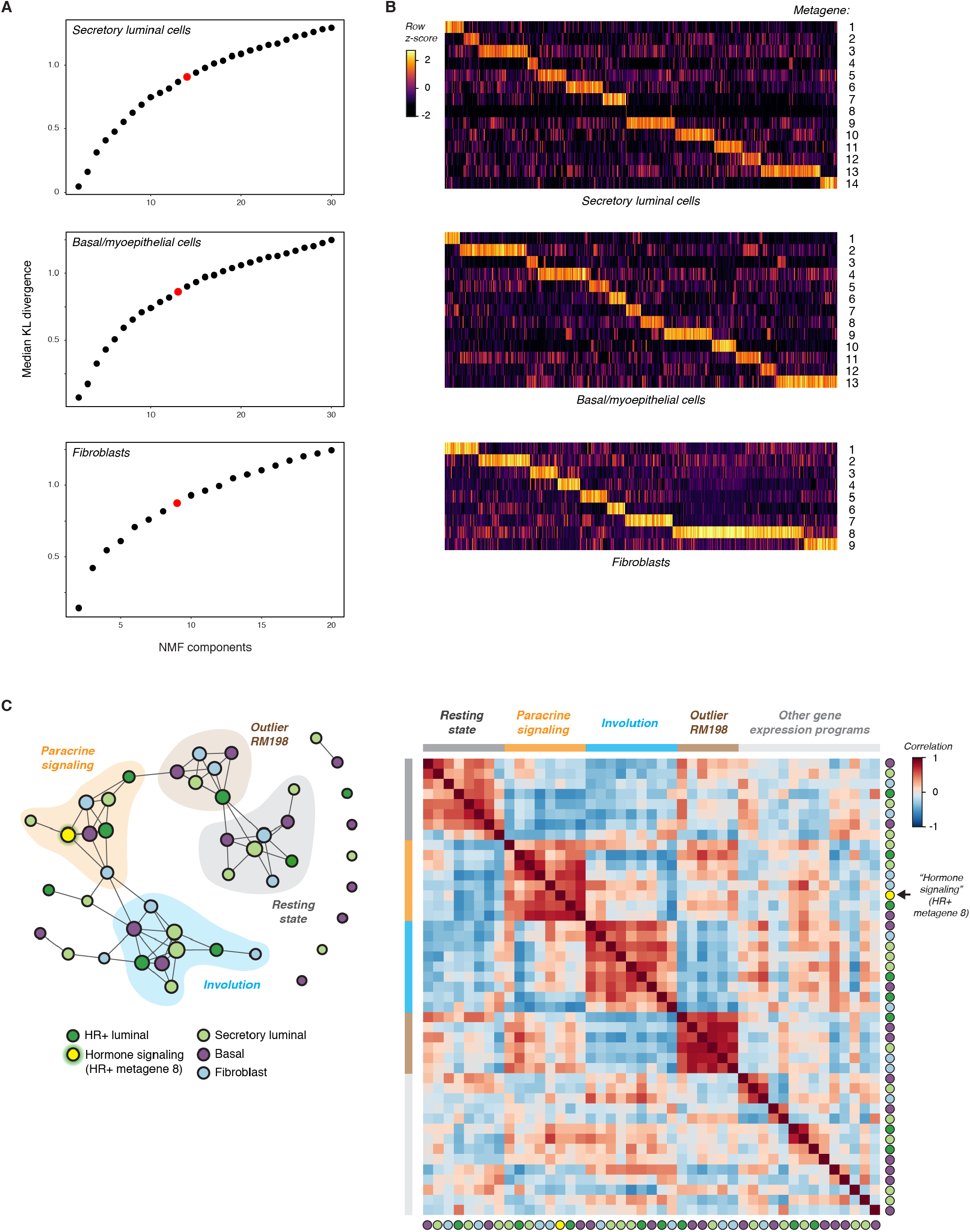
Matrix decomposition analysis of secretory luminal cells, basal/myoepithelial cells, and fibroblasts. (A) Parameter selection for non-negative matrix factorization based on KL divergence (*methods*). (B) Heatmap of cell loadings across each metagene for the indicated cell types. (C) *Left:* Network graph of coordinated gene expression programs in the human breast. Nodes represent distinct metagenes in the indicated cell types, and edges connect highly correlated metagenes (Pearson correlation coefficient > 0.5 and p < 0.05). Modules of highly correlated gene expression programs were identified using the infomap community detection algorithm. The “hormone signaling” metagene in HR+ cells (HR+ metagene 8) is highlighted in yellow. *Right:* Heatmap depicting Pearson correlation coefficients between all metagenes.

**Fig. S13.**
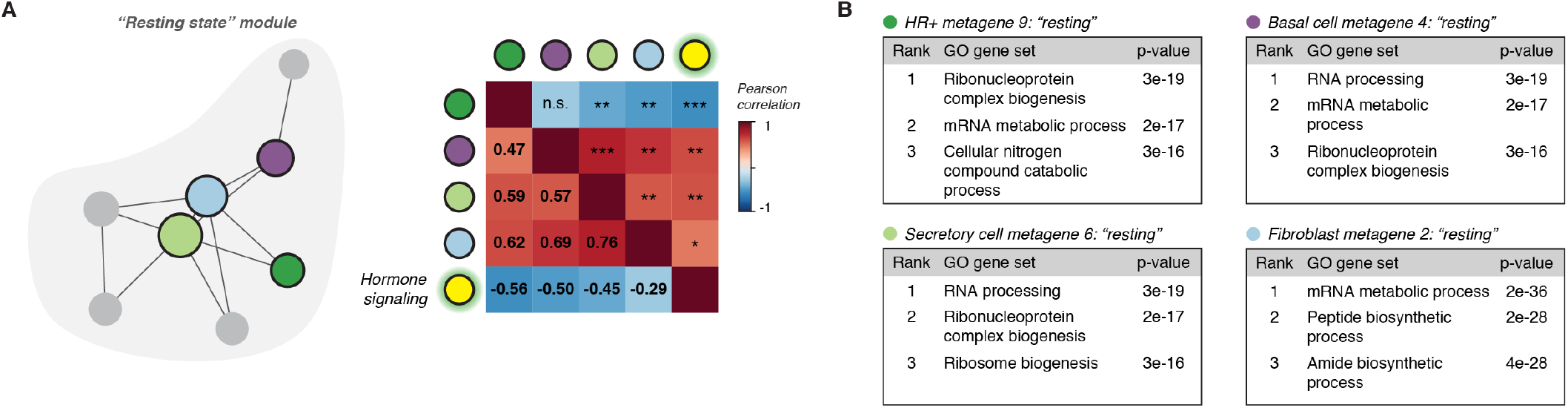
The “Resting State” module consists of metagenes negatively correlated with hormone signaling in HR+ luminal cells. (A) Network subgraph of the “resting state” module, and heatmap depicting Pearson correlation coefficients between the indicated metagenes and levels of significance (* p < 0.05, ** p < 0.01, *** p < 0.001). (B) Gene set enrichment analysis for the indicated metagenes, showing the top pathways identified from GO gene sets.

**Fig. S14.**
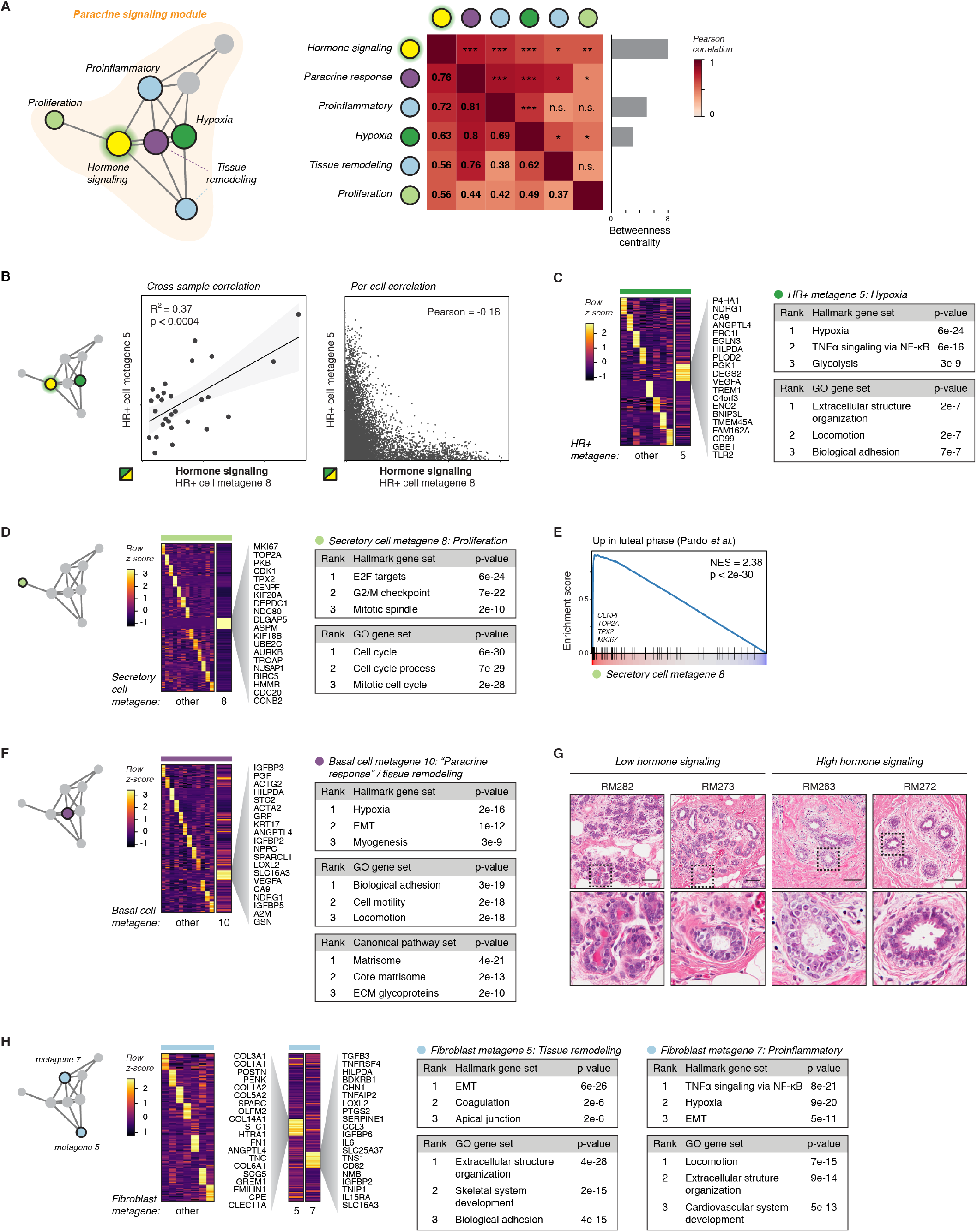
The “Paracrine Signaling” module consists of metagenes positively correlated with hormone signaling in HR+ luminal cells. (A) Network subgraph of the “paracrine signaling” module, and heatmap depicting Pearson correlation coefficients between the indicated metagenes and levels of significance (* p < 0.05, ** p < 0.01, *** p < 0.001). *Right:* Betweenness centrality for the indicated metagenes within the paracrine signaling module. (B) *Left*: Linear regression analysis of HR+ cell state across metagene 8 (“hormone signaling”) versus metagene 5 (“hypoxia”) (R^2^ = 0.37, p < 0.0004, Wald test). Dots represent the average expression of each metagene within a sample. *Right:* Scatter plot of HR+ cell expression of metagene 8 versus metagene 5. Dots represent the expression of each metagene within individual HR+ luminal cells. (C) *Left:* Heatmap depicting the top 20 genes expressed in each HR+ cell metagene, highlighting metagene 5. *Right:* Gene set enrichment analysis for HR+ metagene 5, showing the top pathways identified from the Molecular Signatures Database Hallmark and GO gene sets. (D) *Left:* Heatmap depicting the top 20 genes expressed in each secretory luminal cell metagene, highlighting metagene 8. *Right:* Gene set enrichment analysis for secretory cell metagene 8, showing the top pathways identified from the Molecular Signatures Database Hallmark and GO gene sets. (E) Gene set enrichment analysis of secretory luminal cell metagene 8, showing enrichment of genes upregulated during the luteal phase of the menstrual cycle (NES = 2.38, p < 2e-30) (*29*). (F) *Left:* Heatmap depicting the top 20 genes expressed in each basal cell metagene, highlighting metagene 10. *Right:* Gene set enrichment analysis for basal cell metagene 10, showing the top pathways identified from the Molecular Signatures Database Hallmark, GO, and Canonical Pathways gene sets. (G) Representative images of H&E stained sections. Scale bars 100 µm. (H) *Left:* Heatmap depicting the top 20 genes expressed in each fibroblast metagene, highlighting metagenes 5 and 7. *Right:* Gene set enrichment analysis for fibroblast metagenes 5 and 7, showing the top pathways identified from the Molecular Signatures Database Hallmark and GO gene sets.

**Fig. S15.**
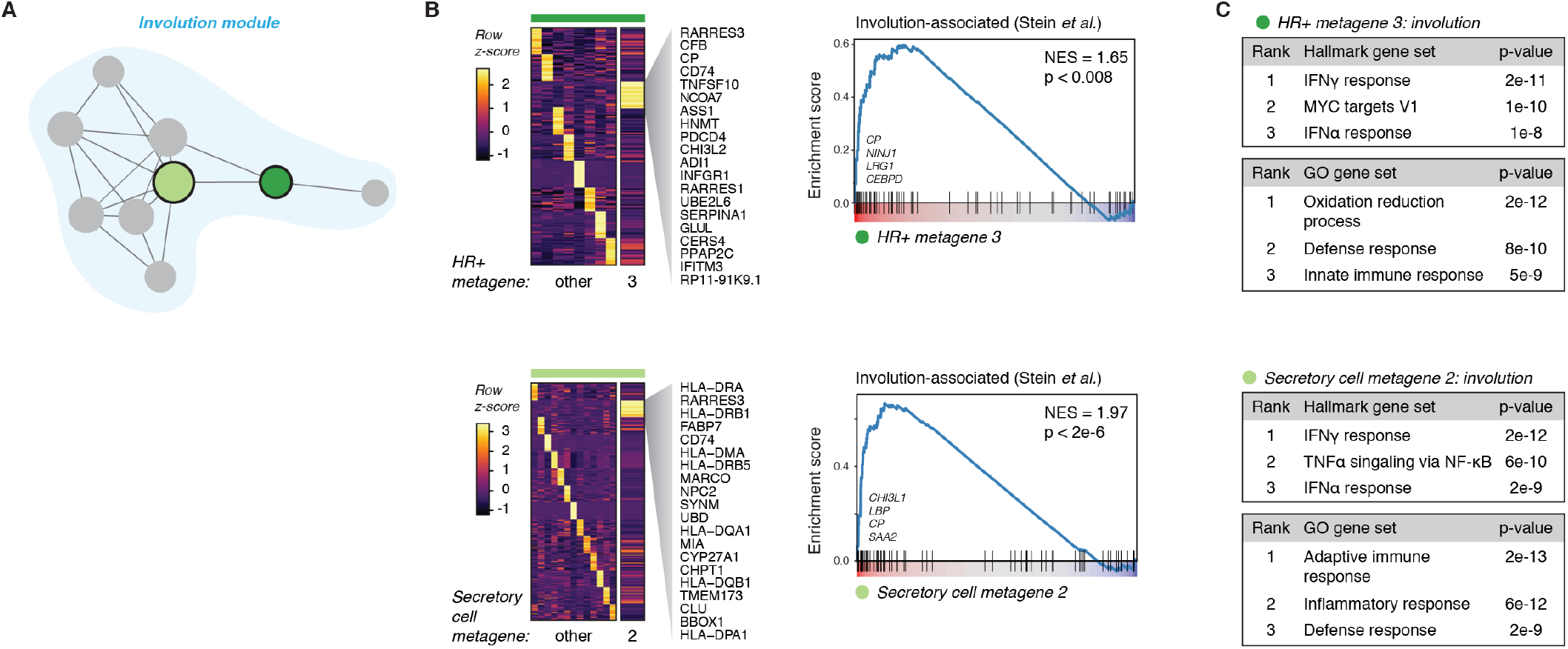
The “Involution” module consists of metagenes enriched for genes upregulated during post-lactational involution. (A) Network subgraph of the “involution” module, highlighting the two metagenes most closely associated with an “involution-like” gene signature. (B) Heatmap depicting the top 20 genes expressed in each HR+ cell or secretory cell metagene, highlighting “involution-like” metagenes. *Right:* Gene set enrichment analysis of the indicated metagenes, showing enrichment of genes upregulated during the postlactational involution (HR+ metagene 3: NES = 1.65, p < 0.008; Secretory cell metagene 2: NES = 1.97, p < 2e-6) (*51*). (C) Gene set enrichment analysis for the indicated metagenes, showing the top pathways identified from the Molecular Signatures Database Hallmark and GO gene sets.

**Fig. S16.**
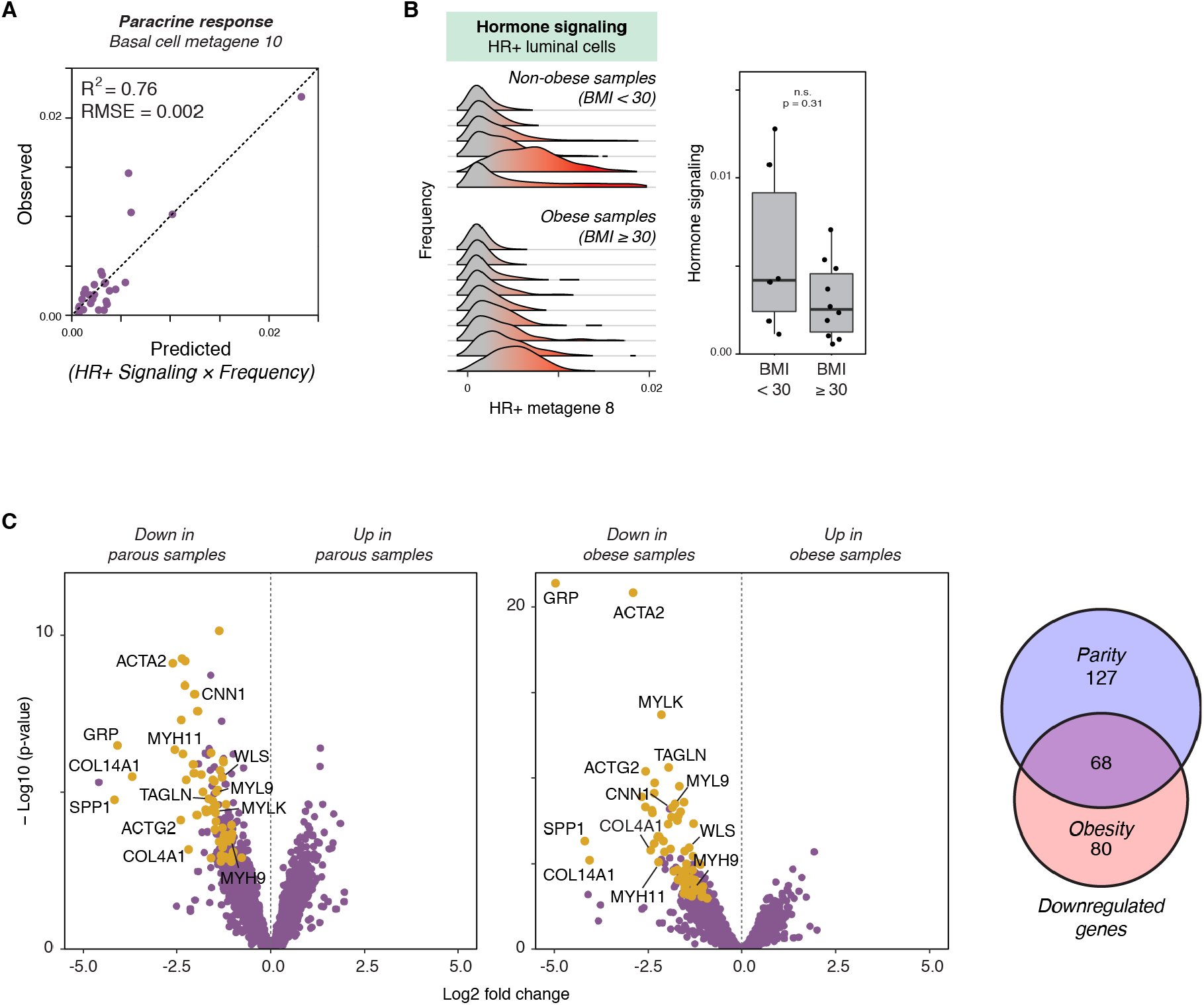
Paracrine signaling to basal cells depends on the hormone signaling state of HR+ luminal cells and the proportion of HR+ luminal cells in the epithelium. (A) Plot depicting the observed basal cell state across metagene 10 (“paracrine response”) for each sample versus the predicted values based on multiple linear regression analysis with three predictors: HR+ cell hormone signaling (HR+ metagene 8), the frequency of HR+ cells in the epithelium, and an interaction term representing the combined effects of HR+ cell signaling and frequency (Signaling × Frequency). (B) Ridge plots depicting the distribution of HR+ cell metagene 8 (“hormone signaling”) expression across samples, and quantification of the average expression in obese (BMI ≥ 30) versus non-obese (BMI < 30) samples (n = 16 samples, p < 0.31, Mann-Whitney test). (C) Volcano plot and venn diagram highlighting genes downregulated in basal/myoepithelial cells in both parous and obese samples.

